# Mapping the endothelial O-GlcNAcome uncovers CCAR1 as a regulator of senescence

**DOI:** 10.64898/2026.02.12.705616

**Authors:** Andreas Will, Regine Heller, Claudia Ender, Felix R. Schneidmadel, Florian Meier-Rosar, Darya Zibrova

## Abstract

O-GlcNAcylation, the reversible addition of O-linked N-acetylglucosamine (O-GlcNAc) to serine and threonine residues, is a dynamic posttranslational modification that integrates nutrient and stress signals to fine-tune protein function and maintain cellular homeostasis. Although dysregulation of O-GlcNAc signaling is associated with age-related pathologies, its role in physiological aging remains unclear. Here we show that chronologically aged human vascular endothelial cells exhibit reduced O-GlcNAcylation due to altered abundance of enzymes in the hexosamine biosynthesis pathway and O-GlcNAc cycling. Conversely, the attenuation of O-GlcNAcylation achieved by genetic or pharmacological means, induced senescence via canonical p53/p21^CIP1^ and p16^INK4a^/Rb pathways and impaired key endothelial functions, such as proliferation and angiogenic capacity. Importantly, O-GlcNAcylated substrates identified across the entire proteome in endothelial cells closely aligned with these mechanistic and phenotypic findings. These substrates were enriched in regulators of processes central to senescence and vascular aging, such as genome stability, transcription, cell cycle, and methylation. Among these, we confirmed cell division cycle and apoptosis regulator 1 (CCAR1) as a bona fide O-GlcNAc substrate and demonstrate that its expression and O-GlcNAcylation decline during senescence in endothelial cells. CCAR1 depletion suppressed apoptosis and sensitized cells to oxidative stress-induced DNA damage, whereas restoration of CCAR1 levels mitigated premature and replicative senescence. Together, our findings establish O-GlcNAc signaling as a key regulatory system that integrates metabolic and stress cues to coordinate protein networks controlling genome stability, transcription and cellular stress adaptation, thereby preserving endothelial integrity and restraining senescence onset.

## Introduction

O-GlcNAcylation is a dynamic post-translational modification of proteins in which a single N-acetylglucosamine (GlcNAc) moiety is attached to serine or threonine residues via a β-glycosidic bond to form O-linked β-N-acetylglucosamine (O-GlcNAc). Similar to phosphorylation, this modification modulates protein function and is tightly regulated by just two enzymes: O-GlcNAc transferase (OGT), which adds O-GlcNAc to proteins, and O-GlcNAcase (OGA), which removes it. Predominantly nucleocytoplasmic, O-GlcNAcylation operates as a key signaling mechanism that orchestrates diverse cellular processes, including transcription, translation and cell division (reviewed in (1)). The donor substrate used by OGT, uridine diphosphate N-acetylglucosamine (UDP-GlcNAc), is generated through the hexosamine biosynthesis pathway (HBP), which is rate-limited by glutamine:fructose-6-phosphate amidotransferase (GFAT) (2). The HBP integrates metabolites derived from glucose, amino acids, fatty acids, and nucleotides into the synthesis of UDP-GlcNAc. This couples O-GlcNAcylation to nutrient and energy availability, thereby fine-tuning protein function according to the metabolic state of the cell (3).

In addition to metabolic cues, the HBP is highly sensitive to various stressors, including heat, high salt levels, heavy metals, UV light, and hypoxia. These stressors rapidly elevate the O-GlcNAcylation of many proteins as a protective response to acute stress (4–8). By contrast, chronic dysregulation of O-GlcNAcylation has been linked to various age-related pathologies, including neurodegeneration, cancer, and, in particular, cardiovascular complications arising in the course of metabolic disorders such as type 2 diabetes (9–12). With regard to the latter, a causal role of altered O-GlcNAc signaling in disease pathogenesis has been established (reviewed in (12)). Mechanistic studies in diabetes models have revealed that hyperglycaemia-induced endothelial dysfunction, an initiating event in the development of vascular complications, is partly mediated by excessive O-GlcNAcylation of key signaling and metabolic regulators, most notably endothelial nitric oxide synthase (eNOS) (13–17).

While these studies have elucidated the pathophysiological mechanisms involving O-GlcNAc under metabolic stress, little is known about how O-GlcNAcylation is regulated during normal vascular aging, where alterations may precede overt disease. Studies in rodents have revealed significant age-dependent changes of O-GlcNAcylation in the cardiovascular system. For example, reduced O-GlcNAc levels have been reported in rat hearts between 6 and 22 months of age coinciding with lower OGT expression (18). Conversely, elevated O-GlcNAc levels have been found in the heart and aorta of 24-month-old rats despite lower OGT abundance (19). However, it is difficult to distinguish whether the observed O-GlcNAc changes originate within cardiovascular tissue or are secondary to metabolic dysregulation in other organs that trigger O-GlcNAc as a systemic nutrient sensor, as animal studies can be confounded by systemic metabolic alterations that accompany advanced age (19).

A hallmark and one of the major underlying mechanisms of vascular ageing and age-related vasculopathies is endothelial senescence, a state of stable cell cycle arrest (20–22). Endothelial cells are particularly susceptible to senescence triggers, such as oxidative stress, chromatin disruption, and DNA damage. These triggers lead to the sustained activation of the p16^INK4a^/Rb and p53/p21^CIP1^ pathways, which inhibit the cyclin-dependent kinases (CDKs) that control the cell cycle (23, 24). Senescent endothelial cells accumulate during normal ageing and are further enriched in pathological contexts such as atherosclerosis and hypertension (25–29). While there is limited direct evidence linking O-GlcNAcylation to senescence, several studies have suggested a mechanistic connection. O-GlcNAcylation regulates the cell cycle progression (30–34), and numerous proteins involved in the control of proliferation. For instance, p53, c-Myc, β-catenin, cyclin D, checkpoint kinases, and mitotic spindle components, are known O-GlcNAc targets (reviewed in (35)). Chemotherapy-induced senescence in cancer cells also alters the HBP and O-GlcNAcylation (36, 37). Taken together, these findings suggest that O-GlcNAcylation may play a pivotal role in coordinating senescence-associated processes. However, most of this evidence originates from immortalized or cancer cell lines, in which O-GlcNAc signaling and cell cycle control can differ markedly from those in primary cells.

In this study, we sought to systematically map the landscape of protein O-GlcNAcylation in primary vascular endothelial cells by employing an antibody-based O-GlcNAc enrichment (38) and trapped ion mobility–mass spectrometry (TIMS-MS) (39). By applying this technology alongside cellular senescence models, we gained mechanistic insight into the interplay between O-GlcNAcylation, senescence, and endothelial dysfunction. This opens up potential new avenues for therapeutic interventions in age-related vascular disorders. As a potential target, we identified cell division cycle and apoptosis regulator 1 (CCAR1) as an O-GlcNAc substrate, which is involved in the regulation of premature and replicative senescence in endothelial cells.

## Materials and methods

### Chemicals

Cell culture media (M199) was from Lonza. Fetal calf serum, human serum, endothelial growth factor supplement were purchased from Sigma-Aldrich. Human serum albumin was from Bayer Vital. Recombinant human VEGF-165 was obtained from R&D Systems, Inc. Human plasma fibrinogen was from Merck Millipore. EDTA-free protease inhibitor mixture complete and Cell Proliferation ELISA (BrdU, colorimetric, #11647229001) were purchased from Roche. DC^TM^ Protein Assay kit was acquired from Bio-Rad Laboratories and Pierce BCA Protein Assay Kit from Thermo Scientific (#23227). Western Lightning^®^ Plus-ECL (#NEL105001EA) was obtained from Revvity. Protein G Sepharose (#17061801) was from Cytiva. Senescence β-Galactosidase Staining Kit (#9860) was from Cell Signaling Technology. SAINT-RED was purchased from Synvolux Therapeutics B.V. Trypsin was obtained from Sigma-Aldrich (#T6567), Lys-C from Wako Chemicals (#125-05061) and Chymotrypsin from Promega (#V1061). C18 extraction disks (2215-C18) were from Thermo Scientific (#66883-U). ReproSil-Pur C18-AQ (1.9 µm) resin was purchased from Dr. Maisch GmbH. The PTMScan® O-GlcNAc [GlcNAc-S/T] Motif Kit was from Cell Signaling Technology (#95220S). All other reagents and solvents were from Sigma-Aldrich unless indicated otherwise.

### Antibodies

The following antibodies were purchased from Cell Signalling Technology: O-GlcNAc (CTD110.6 clone, #9875), OGT (#24083), p16^INK4A^ (#92803), p21^Cip1^ (#2947), phospho(T202/Y204)-Erk1/2 (#9106), Erk1/2 (#9107), phospho(S139)-H2A.X (#9718), phospho(S15)-p53 (#92845), phospho(S46)-p53 (#2521), phospho(S780)-Rb (#9307), β-actin (#4970); caspase-3 (#9665), cleaved caspase-3 (#9664), PTMScan^®^ O-GlcNAc [GlcNAc-S/T] Motif Kit (#95220). O-GlcNAc antibody (RL2 clone, #677902) was from BioLegend. OGA (#sc-376429) and p53 (#sc-126) antibodies were purchased from Santa Cruz Biotechnology. CDK2 antibody (#ab32147) was from Abcam. CCAR1 antibody (#A300435A) was obtained from Bethyl. Polyclonal sheep antibody against GFAT1 (#S702C, 3rd bleed) were generated in the Division of Signal Transduction Therapy (University of Dundee, Scotland, UK) by immunization with a full-length human GST-GFAT1. Horseradish peroxidase (HRP)-conjugated secondary antibody to rabbit and mouse IgG were from Kirkegaard & Perry Laboratories, Inc. (KPL). HRP-conjugated secondary antibody to sheep IgG (#sc-2056) was from Santa Cruz Biotechnology. HRP-conjugated secondary antibody to mouse IgM µ chain (#074-1803) was purchased from KPL.

### Primary endothelial cells, senescence models and treatment conditions

Human umbilical vein endothelial cells (HUVEC) were isolated from anonymized umbilical cords in accordance with the Declaration of Helsinki (1964). Collection and use of the tissue were approved by the ethics committee of Jena University Hospital (2023-2894-Material), and written informed consent was obtained from all donors. The study included data generated from around 50 independent HUVEC isolations, with 3–5 distinct donor-derived batches used per experimental series. In general, cells of the first or second passage were seeded for the experiments implying treatment with siRNA, lentiviruses or chemical compounds.

HUVEC, isolated by collagenase treatment, were maintained in M199 supplemented with 17.5% fetal calf serum (FCS), 2.5% human serum, 100 U/mL penicillin, 100 µg/mL streptomycin, 680 µM glutamine, 25 µg/mL heparin, 7.5 µg/mL endothelial cell growth supplement and 100 µM vitamin C (growth medium), following established protocols (40). To induce premature senescence, HUVEC were seeded at a density of 23,000 cells/cm² on collagen-coated dishes and, after 48 h, were exposed to H₂O₂ (100 µM) for 8 days with the H₂O₂-containing medium replaced daily. Control cells were handled identically but received H₂O₂-free medium. After 8 days, the cells were harvested either for biochemical analysis or used for senescence-associated β-galactosidase (SA-β-gal) staining. Replicative senescence was induced by serially passaging primary HUVEC from passage 1 (P1) to later passages (P20, P25 or P30). During early passages, cells were split at a 1:4 ratio and later, when the proliferation slowed down, at a 1:2 ratio. Passaging was performed at around 80% confluency to prevent growth arrest by contact inhibition. At the end of each senescence protocol, the cells were harvested for biochemical analyses or subjected to SA-β-galactosidase staining.

To modulate O-GlcNAc levels pharmacologically, HUVEC monolayers were treated either with the OGA inhibitor Thiamet G (2 µM, 3 h) or the OGT inhibitor OSMI-1 (50 µM, 4 days) with cells treated with the diluent DMSO serving as control.

### SA-β-gal staining

SA-β-gal activity was assessed using the Senescence β-Galactosidase Staining Kit according to the manufacturer’s instructions. Briefly, the cells were seeded at a density of 5,555 cells/cm^2^ on gelatin-coated 35 mm-diameter dishes. After 48 h, the cells were washed once with PBS and fixed with 1 mL of fixative solution for 15 min at room temperature. Following two PBS washes, 1 mL of staining solution was added. The dishes were then sealed with parafilm and incubated at 37 °C in a dry incubator for up to 24 h. Afterwards, the staining solution was removed and the cells were overlaid with 70% glycerol. Images were acquired using an AxioVert 200 light microscope (Carl Zeiss). Quantification was performed using cellSens™ software (Olympus) whereby SA-β-gal-positive cells were scored relative to the total number of cells. Five to ten random fields, each containing between 30-300 cells/field, were analyzed per condition.

### Cell lysis and protein-immunoprecipitation

After the respective treatments, cells were rinsed with PBS and lysed on ice in buffer containing 50 mM Tris (pH 7.5), 1 mM EDTA, 1 mM EGTA, 1% (v/v) Triton X-100, 1 mM Na₃VO₄, 50 mM NaF, 5 mM Na₄P₂O₇, 0.27 M sucrose, 0.1% (v/v) β-mercaptoethanol, 0.2 mM PMSF, 1% complete protease inhibitor cocktail, 40 µM PUGNAc and 2 µM Thiamet G (the last two components were added for preservation of O-GlcNAcylation). Lysates were clarified by centrifugation (13,000 × g, 10 min, 4°C), and protein concentrations were determined using DC™ Protein Assay kit (Bio-Rad) according to Lowry method, with bovine serum albumin used as the standard.

For immunoprecipitation (IP) of endogenous CCAR1, lysates (0.3-0.5 mg protein in 500 µl of lysis buffer) were incubated overnight at 4°C on a rotating wheel with 2.5 µg of CCAR1 antibody and 10 µl of 50% Protein G Sepharose. The Protein G Sepharose beads carrying the immune complexes were then collected by centrifugation at 6,000 × g for 1 min, after which they were washed sequentially on ice with lysis buffer containing 0.5 M NaCl, salt-free lysis buffer, and 50 mM Tris (pH 8.0) containing 0.1 mM EGTA. Two washes per condition were performed. The bound proteins were eluted by boiling the beads in 25 µL/pellet of 2× Laemmli buffer at 95°C for 5 min and then recovered by centrifugation at 16,000 × g for 5 min at room temperature. This step was repeated twice. The resulting supernatants were subjected to immunoblotting. For CCAR1 interactome analysis, the IP was performed essentially as described above except that the immunoprecipitates were eluted with 25 µL of 0.15 % trifluoroacetic acid (TFA) at room temperature for 10 min. A total of 3 elutions were pooled. The pH of the eluate was neutralized by the addition of 1 M Tris (pH 10) at a ratio of 1:10 and the sample was further analyzed by MS.

### Immunoblotting

Cell lysates (10-30 µg protein per lane) or immunoprecipitates were resolved by SDS-PAGE and transferred to PVDF membranes. The membranes were blocked for 1 h in TBST (20 mM Tris-HCl (pH 7.6), 137 mM NaCl, 0.1% (v/v) Tween-20) supplemented with 5% non-fat dry milk or with 4% BSA (for O-GlcNAc detection). Primary antibodies were then applied overnight at 4°C in TBST containing 5% BSA. Following a 1-h incubation with HRP-conjugated secondary antibodies at room temperature, the signals were visualized using enhanced chemiluminescence. Band intensities were quantified by densitometry using ImageJ software and the values were normalized to the corresponding loading control or, when applicable, phospho-to-total protein ratios were calculated. To estimate global O-GlcNAc levels, all detectable O-GlcNAc-reactive bands within a lane were quantified and summed for each condition. These cumulative values were then normalized to the corresponding loading controls and compared across treatments.

### RNA interference-mediated gene knockdown

Small interfering RNA (siRNA) duplexes targeting human GFAT1, OGT and CCAR1 were used. SMARTpool siRNAs against OGT and CCAR1 (M-019111-00-0005, M-013828-00-0005, respectively) and a non-targeting SMARTpool control (D-001810-10-20) were obtained from Dharmacon/GE Healthcare. For GFAT1, the duplex reported by Jokela et al. (41) (sense: 5′-GGAGGAUACUGAGACCAUU-3′; antisense: 5′-U UACCAGU CAGUAUCCUCC-3′) was purchased from Sigma-Aldrich. HUVEC were seeded 24 h before transfection at a density of 18,000 cells/cm² onto gelatin-coated 100 mm-diameter dishes. The cells reached approximately 50% confluence at the time of transfection. A total of 4 µg siRNA per dish was complexed with 60 nmol of the SAINT-RED transfection reagent (Synvolux Therapeutics) and diluted in M199 containing 0.25% of human serum albumin to a final volume of 4 ml. The complexes were incubated for 4 h, after which 8 ml of growth medium was added. The cells were then cultured for 72 h, after which they were reseeded and subjected to a second transfection cycle for an additional 96 h following the same procedure. After the second cycle was complete, the cells were harvested for biochemical analyses or used for BrdU incorporation or spheroid assays.

### Lentiviral transduction for recombinant CCAR1 overexpression

Lentiviral particles were produced in HEK293T cells to enable stable CCAR1 overexpression in HUVEC. 24 h before transfection, HEK293T cells were seeded at a density of 18,200 cells/cm^2^ on 100 mm-diameter dishes and co-transfected with an mGFP-tagged human CCAR1 expression vector at a concentration of 10 µg/dish (#RC224293L4, OriGene) as well as with the following packaging plasmids: pMDL (10 µg/dish), pRSV (5 µg/dish) and pVSV-G (2 µg/dish) kindly provided by Dr. Jörg Müller (Jena University Hospital, Germany). Transfection was performed using PEI (67.5 µL of 1 mg/mL PEI per dish) and mGFP fluorescence was assessed 24 h later. Typical transfection efficiencies were greater than 80%.

Viral supernatants were collected at 24, 48 and 72 h. They were then cleared by filtration (0.22-µm low-protein binding filter) and concentrated using Amicon^®^ Ultra centrifugal filters (30 kDa cut-off; 4,000 × g, 15 min, room temperature). After each harvest, the HEK293T cultures were replenished with fresh medium. The concentrates of viral particles recovered from 6 × 90-mm dishes at each time point were distributed across all the wells of a 12-well plate containing HUVEC that were seeded 24 h earlier (40,000 cells/well on double gelatine-coated surfaces). Then, the cells were spinfected (500 × g, 1 h, room temperature) in the presence of 8 µg/mL polybrene. Puromycin (0.5 µg/mL) was added 96 h after the final infection to select stable transductants. Cultures that reached confluence within approximately one week were expanded and used for downstream experiments. Untransduced HUVEC from the same donor were handled in parallel with the CCAR1-transduced cultures and underwent the same processes, including exposure to polybrene and centrifugation during mock spinfection. These cells served as control cells.

### BrdU incorporation assay

HUVEC were seeded at a density of 3,500 cells per well on 96-well plates and allowed to attach overnight in growth medium. The cells were then starved for 4 h in low-serum M199 (2% FCS; identical supplements as in complete growth medium except human serum and endothelial mitogen) and subsequently stimulated with basic fibroblast growth factor (bFGF, 50 ng/ml) for 24 h.

BrdU incorporation was quantified using a colorimetric ELISA kit according to the manufacturer’s instructions. Briefly, 3 h before the end of the 24 h stimulation period with bFGF, the BrdU labelling solution was added to achieve a final concentration of 10 µM. After the medium was removed, the cells were fixed and the DNA was denatured. Then, an anti-BrdU peroxidase-conjugated antibody was applied for 90 min. Following PBS washes, substrate solution was added and the reaction was stopped after 15 min with 1 M H_2_SO_4_. Absorbance was measured at 450 nm using a microplate reader.

### Spheroid assay

Spheroids were generated as previously described (42). HUVEC with reduced O-GlcNAcylation, achieved through combined GFAT1/OGT knockdown or OSMI-1 treatment, were resuspended in growth medium and mixed at a 5:1 ratio with methylcellulose (12 mg/mL). Subsequently, 3,000 cells per well were seeded into 96-well round-bottom plates and cultured for 24 h to allow spheroid formation. The spheroids were then collected, washed with HEPES buffer, and transferred to a fibrinogen solution (1.8 mg/mL fibrinogen, 20 U/mL aprotinin). This suspension containing approximately 100 spheroids/mL was mixed with thrombin (0.2 U/well) before being added to 24-well plates (300 µL/well). The plates were then incubated at 37°C for 20 minutes to allow polymerization into three-dimensional fibrin gels. Thereafter, thrombin was removed by washing the gels with M199 containing 2% FCS and the gels were overlaid with the same medium. Sprouting was induced by addition of 10 ng/ml vascular endothelial growth factor (VEGF) for 24 h. The spheroids were then fixed with 4% paraformaldehyde, imaged by bright-field microscopy (AxioVert 200, Carl Zeiss) and quantified using the cellSens™ software (Olympus). For each experiment, at least ten spheroids per condition were analyzed for sprout number, mean sprout length and cumulative sprout length.

### Statistical analysis of biochemical data

The data are presented as mean ± SEM from 3-5 independent experiments. Comparisons between two groups were performed using a Student’s t-test. For datasets involving more than two groups, one-way ANOVA with Tukey’s post hoc test was used or two-way repeated measures ANOVA with Bonferroni correction where applicable. Statistical significance was defined as p<0.05. All analyses and graphing were performed using GraphPad Prism 8.0.2.

### O-GlcNAc enrichment at the peptide level

Proteome-wide enrichment of O-GlcNAcylated peptides was performed using the PTMScan® O-GlcNAc [GlcNAc-S/T] Motif Kit (Cell Signaling Technology). The cells were seeded on 90 mm culture dishes and cultured for 3 days until they reached confluency. They were then treated with 2 µM Thiamet G for 3 h rinsed with PBS and lysed on ice in buffer containing 20 mM HEPES, 6 M guanidinium hydrochloride, 2.5 mM sodium pyrophosphate, 1 mM β-glycerophosphate, 1 mM sodium orthovanadate, and 2 µM Thiamet G. The samples were sonicated using a tip sonicator, snap-frozen in liquid nitrogen and stored at −20 °C. After thawing, the lysates were centrifuged (20,000 × g, 10 min) and the pellets were discarded. The protein concentrations in the supernatant were determined using the Pierce BCA Protein Assay Kit.

For each biological replicate, we processed 4 mg of protein. The samples were reduced and alkylated with TCEP (10 mM) and CAA (40 mM) (7 min, 45 °C, 1,000 rpm). They were then diluted with 50 mM HEPES (pH 8.5) to a final guanidinium concentration <0.5 M, and digested overnight with Lys-C (1:200) and trypsin (1:100) at 37 °C with shaking (800 rpm).

The digested samples corresponding to 4 mg protein were combined and desalted using Sep-Pak tC18 cartridges (500 mg sorbent). The columns were then equilibrated with acetonitrile, Sep-Pak Buffer A (0.1% formic acid, 50% acetonitrile), and Sep-PakB B (0.1% TFA). The samples were acidified to 1% formic acid, incubated on ice for 15 min, and then centrifuged (20,000 × g, 5 min, 4 °C). The resulting supernatants were loaded onto the columns. After washing, the peptides were eluted with 1.8 mL Buffer A, snap-frozen, and lyophilized overnight (VaCo 2, Zirbus, Bad Grund, Germany).

Immunoprecipitation of O-GlcNAcylated peptides was performed according to the manufacturer’s instructions using one-third of the bead volume provided per vial and 1 mg peptide input per sample, as determined by absorbance at 280 nm. Eluates were desalted using C18 StageTips.

### C18 StageTip desalting and purification

Prior to LC-MS analysis, the peptide samples were desalted and purified using C18 stage tips (43), prepared with two layers of Empore 2215-C18 Octadecyl 47 mm disks (Thermo Scientific). Buffers were added in 50-µL volumes with centrifugation at 300 × g. The stage tips were equilibrated with 100% ACN, followed by one volume of Buffer A (0.1% TFA, 40% ACN), and two volumes of Buffer B (0.1% TFA). Acidified samples (pH < 4) were applied to the StageTip followed by two washes (Buffer B) and subsequent elution (Buffer A). The samples were then dried in a vacuum centrifuge (45 °C, 40 min) and reconstituted in MS loading buffer for timsTOF HT (2% ACN, 0.3% TFA), or timsTOF Ultra (0.015% DDM, 0.1% FA). Peptide concentrations were determined by absorbance at 280 nm.

### Mass Spectrometric Data Acquisition

All solvents were HPLC grade. Nanoflow reversed-phase liquid chromatography was performed on a nanoElute or nanoElute 2 system (Bruker Daltonics). The peptides were separated either on a 25 cm column (protein immunoprecipitation) or a 15 cm column (peptide immunoprecipitation), with a 60 min gradient at a flow rate of 0.5 µL/min at 50 °C. The columns were prepared in-house with a pulled emitter tip, packed with 1.9 μm ReproSil-Pur C18 - AQ beads. Mobile phases A and B were water/0.1% formic acid and acetonitrile/0.1% formic acid. The LC system was connected online to a TIMS quadrupole time-of-flight mass spectrometer (timsTOF HT or timsTOF Ultra) via a Captive Spray nano-electrospray source (44).

MS data acquisition was performed as described previously (45). The dda-PASEF method fragments each precursor with an ion mobility-dependent collision energy ranging linearly from 59 eV at 1.60 V·s/cm^2^ to 20 eV at 0.6 V·s/cm^2^. The collision cell radio frequency (RF) was set to 1,500 V, and the pre-pulse storage time was set to 12 μs with a 60 μs transfer time. To improve the detection of diagnostic fragment ions with m/z < 200, we activated the “stepping” option to acquire each fragment ion spectrum twice, once with the collision parameters set as above, and a second time with a reduced collision cell RF of 800 V, a pre-pulse storage time of 8 μs and 25 μs transfer time. During post-processing, both spectra were merged to resemble the standard dda-PASEF data structure.

### Proteomics data analysis

Raw data were searched against the Homo sapiens proteome FASTA (UP000005640_9606_additional, June 16, 2022; 81,131 entries) using FragPipe (v22) with the Glyco-O-HCD workflow (46) with minor adaptations. Contaminants were included and the database was supplemented with 50% reverse decoys. MSFragger search (47) was performed with ‘stricttrypsin’ (Trypsin/P) or alternatively with ‘chymotrypsin’, both with up to 2 missed cleavages. Methionine oxidation and protein N-terminal acetyl were set as variable modifications, and cysteine carbamidomethylation as fixed modification. For quality control, variable modifications of +8.014199 Da (lysine, ^13^C₈) and +10.027228 Da (arginine, ^13^C₁₀) were included to identify spiked-in stable isotope–labeled peptides. Post-processing was performed with Philosopher with 1% false discovery rate (FDR) filtering at the peptide spectral match (PSM) and protein levels (sequential filtering). PTM-Shepherd was enabled with default parameters for open search and “Glyco mass shift” as annotation source. Diagnostic feature extraction was activated. Glycans were assigned with 1% FDR and a mass tolerance of 50 ppm. MS1 quantification was enabled, normalizing intensities across runs and without matching between runs. Generally, the normalized ion intensity was used as the intensity metric. Fragmentation spectra were visualized with the FragPipe-PDV viewer (48).

### Data processing for RL2 immunoprecipitation

For MS analysis of the RL2 immunoprecipitation, the “combined_protein.tsv” was analyzed. Proteins detected in the IgG control were discarded and we stringently required proteins to be detected in all three replicates of either of the two remaining conditions. The individual “PSM.tsv” files were merged and filtered for HexNAc identifications with a glycan FDR < 0.01. GO enrichment was performed using ClusterProfiler (v.4.14.6), utilizing gene annotations from org.Hs.eg.db (v.3.20.0).

### Data processing for O-GlcNAc immunoprecipitation at the peptide level

The “combined_modified_peptide.tsv” was used to visualize the overlap of HexNAc peptide identifications between replicates. Each replicate was filtered for distinct HexNAc peptide sequences, quantified by ion intensity. For protein level information, data was further collapsed to distinct gene names.

### Data processing for CCAR1 co-immunoprecipitation

To visualize the sequence coverage of HexNAc modified peptides on CCAR1, the “combined_ions.tsv” file was filtered for the highest intensity ion for each modified sequence area, excluding ions with alternative cleavage profiles.

To calculate HexNAc peptide fold changes between treatment conditions, the “combined_modified_peptides.tsv” was filtered for the highest intensity entry for each HexNAc-modified peptide sequence (reducing entries with different localization) and the corresponding non-HexNAc entry. Fold-changes between treatment conditions were calculated for each biological replicate separately.

For analysis of CCAR1 co-immunoprecipitation, the “combined_protein.tsv” file was analyzed. To generate volcano plots, the file was analyzed with the Perseus software (v2.1.3) (49). Proteins were filtered for non-human contaminants. Proteins were required to have at least 3 valid values in one of the immunoprecipitation conditions. Missing values were imputed from a normal distribution with Perseus standard conditions (width: 0.3, down-shift: 1.8). Conditions were compared using Student’s T-test with Benjamini-Hochberg FDR for multiple sample testing. The resulting table was exported as a “.txt” file and visualized in RStudio.

### Data availability

The mass spectrometry data underlying this study have been deposited to the ProteomeXchange Consortium (50, 51) via the partner repository with the dataset identifier PXD073426.

## Results

### Cellular senescence reduces global O-GlcNAcylation in endothelial cells

In order to study the effects of cellular senescence on O-GlcNAcylation and its regulatory enzymes, we first induced premature senescence in primary HUVEC by treating them with 100 µM H_2_O_2_ for 8 days (oxidative stress-induced senescence). Senescence was confirmed by an increased proportion of cells positive for SA-β-gal displaying the enlarged morphology typical of senescent cells (up to 98%) (Fig. 1A). This was accompanied by elevated p53 phosphorylation at serine 15 and p21 upregulation (Fig. 1B). Total O-GlcNAc levels, as detected by the CTD110.6 antibody, were reduced 2-fold in senescent cells (Fig. 1C). This effect was paralleled by a reduction in GFAT1 levels, which is the rate-limiting enzyme of the HBP. The levels of OGT and OGA, which control O-GlcNAc cycling, remained unchanged (Figure 1C).

**Figure 1.**
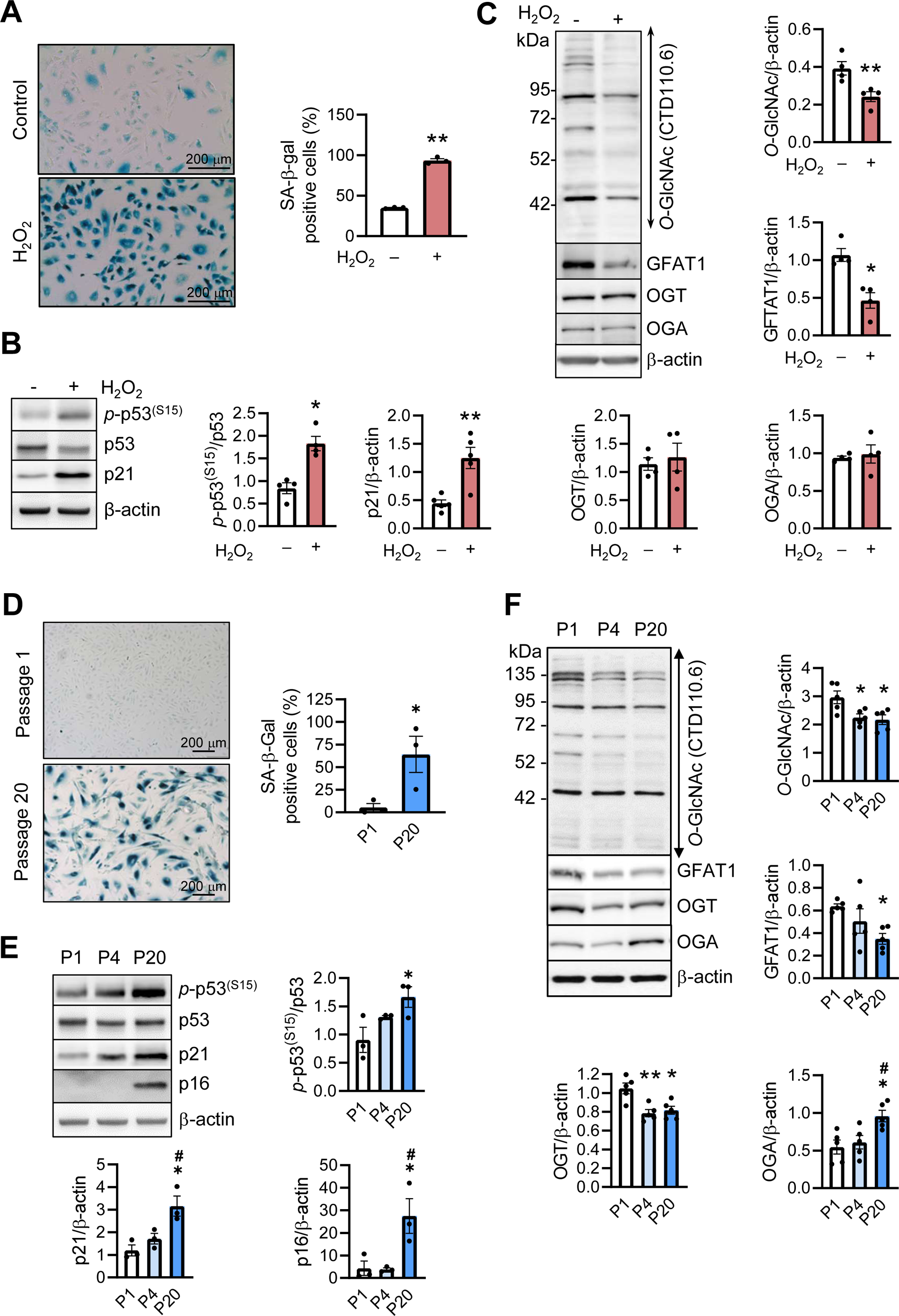
O-GlcNAcylation and its key regulatory enzymes are downregulated in endothelial cells during senescence. **(A-C)** Premature senescence was induced by treating HUVEC with H_2_O_2_ (100 μM, 8 days) with vehicle-treated cells serving as control. **(D-F)** Replicative senescence was achieved by continuous culturing HUVEC up to passage 20 (P20); HUVEC of passage 1 (P1) were analyzed in parallel. In both models, cells were analyzed using SA-β-gal staining or harvested for immunoblotting. Representative immunoblots **(B, C, E, F**) and light microscopy images **(A, D)** with corresponding analyses are shown. Data are means ± SEM of 3–5 independent experiments using cells from different donors. **(A-D)** *p<0.05, **p<0.01 vs. control, paired or unpaired **(D)** t-test. **(E, F)** *p<0.05, **p<0.01 vs. P1; ^#^p<0.05 vs. P4, one-way ANOVA with Tukey’s multiple comparison test.

To confirm these findings in a physiologically relevant context, we turned to a replicative senescence cell model, in which senescent cells are obtained through repeated passaging. This model recapitulates features of chronologically aged endothelial cells (52). At passage 20 (P20), replicative senescence was evident from an increased number of SA-β-gal-positive cells (64% compared to 5% in cells from passage 1 (P1), Fig. 1D). Furthermore, cells at P20 exhibited the typical molecular signatures of replicative senescence including elevated p53 phosphorylation at serine 15, upregulation of p21 and strong induction of the late senescence marker p16 (Fig. 1E). Similar to the premature senescence model above, we found global protein O-GlcNAcylation significantly decreased at both passages 4 and 20 in comparison to cells from P1 (Fig. 1F). This effect was mirrored by a reduction in GFAT1 levels at P20 compared to P1 (Fig. 1F). Furthermore, OGT levels decreased slightly beginning at passage 4 (P4), and OGA levels increased significantly at P20 (Fig. 1F).

Taken together, our data suggest that O-GlcNAcylation is reduced in both prematurely and replicatively senescent endothelial cells, which is most probably the result of an altered abundance of key regulatory enzymes.

### O-GlcNAc depletion induces endothelial cell senescence

To investigate whether the downregulation of O-GlcNAcylation is mechanistically involved in endothelial senescence, we depleted O-GlcNAc in HUVEC by co-transfecting them with GFAT1- and OGT-specific siRNAs, thereby mimicking the conditions observed in replicative senescence. To achieve sustained O-GlcNAc depletion over a prolonged period of time, two consecutive 96-h transfection rounds were performed, resulting in reduced GFAT1 and OGT levels by 84% and 73%, respectively. Consequently, we observed a significant reduction (approximately 3-fold) in total O-GlcNAc levels compared to cells transfected with control siRNA (Fig. 2A). O-GlcNAc-depleted cells showed several characteristics of senescent cells. These included a significant increase in expression of the senescence markers p21 and p16, which both act as CDK inhibitors, a significant decrease in CDK2 levels and a reduced phosphorylation of the retinoblastoma tumor suppressor protein (Rb), which is a master regulator of the cell cycle (Fig. 2A). Furthermore, GFAT1/OGT double knockdown increased the fraction of SA-β-gal positive cells to 25% compared to 5% in control siRNA-treated cells (Fig. 2B). Interestingly, O-GlcNAc depletion stabilized p53, an upstream component of senescence signaling. However, this stabilization was not accompanied by phosphorylation of p53 at serine 15, indicating no activation of classical DNA damage response (DDR) signaling, as supported by unchanged levels of γH2A.X (Suppl. Fig, 1A). Instead, we found p53 increasingly phosphorylated at serine 46, possibly as a consequence of activated Erk signaling (Suppl. Fig. 1B, C).

**Figure 2.**
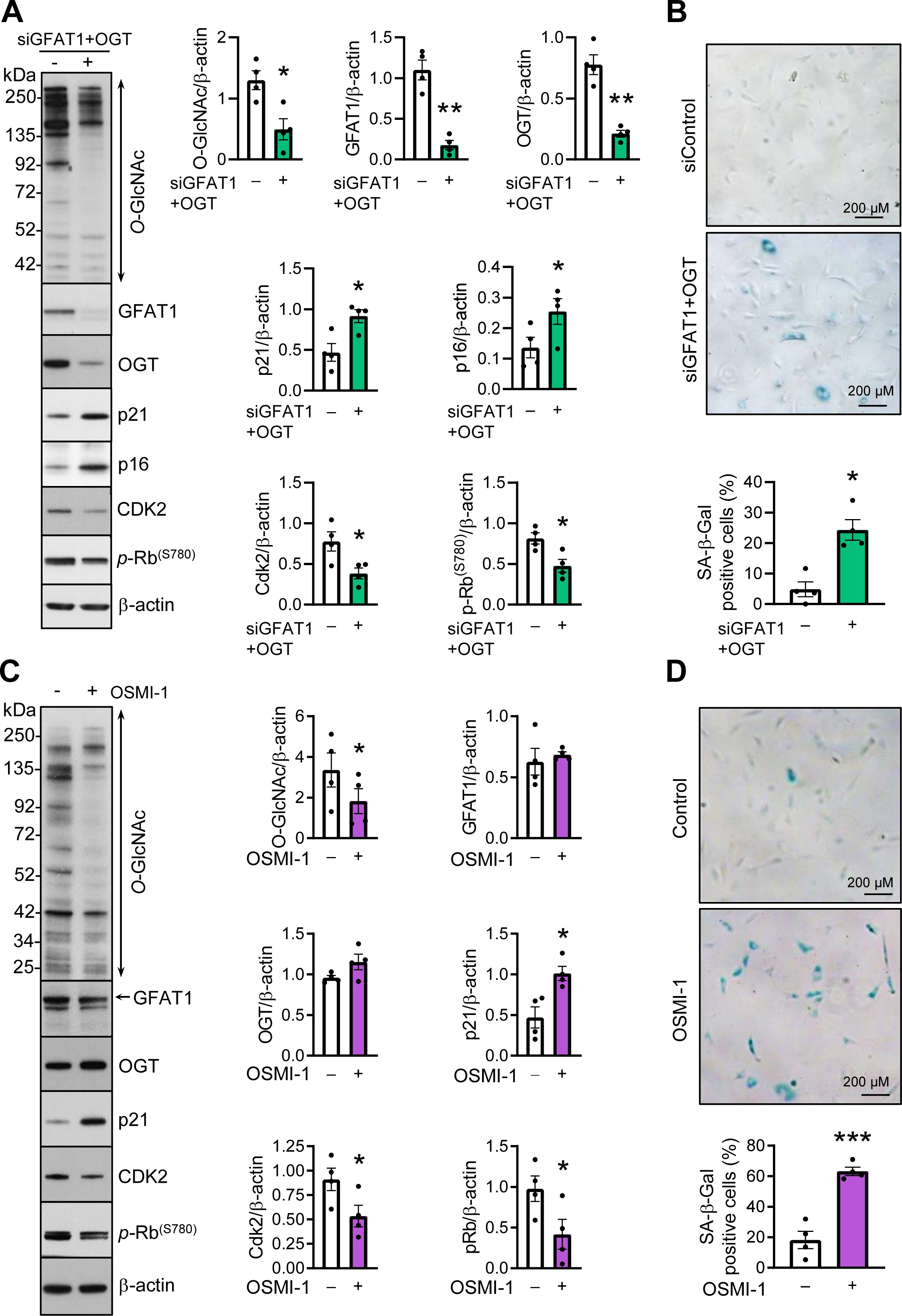
O-GlcNAc depletion induces senescence in endothelial cells. **(A, B)** HUVEC were repeatedly transfected with non-targeting siRNA (-) or co-transfected with GFAT1-and OGT-specific siRNAs (siGFAT1+OGT) (96h, twice). **(C, D)** Cells were treated either with OSMI-1 (50 µM, 96 h) or solvent as control (-). After each intervention, cells were harvested for immunoblotting or analyzed using SA-β-gal staining. Representative immunoblots **(A, C)** and light microscopy images **(B, D)** with corresponding analyses are shown. Data are means ± SEM of 4 independent experiments using cells from different donors. *p<0.05, ***p<0.001 vs. control, paired t-test.

To rule out O-GlcNAc-independent induction of senescence by gene silencing, as an alternative approach to interfering with O-GlcNAcylation, we used OSMI-1, a selective inhibitor of the O-GlcNAc transferase activity of OGT (53). Treatment with OSMI-1 reduced global O-GlcNAcylation 2-fold without markedly affecting GFAT1 or OGT levels (Fig. 2C). Similar to the effects observed in cells with GFAT1/OGT double knockdown, pharmacological depletion of O-GlcNAc levels reproduced the senescence phenotype as demonstrated by increased p21 expression, reduced CDK2 levels and diminished Rb phosphorylation (Fig. 2C). Furthermore, the proportion of SA-β-gal positive cells increased after treatment with OSMI-1as compared to diluent-treated controls (up to 63%)) (Figure 2D).

Together, our findings establish O-GlcNAc depletion as a direct trigger of endothelial cell senescence.

### Disrupted O-GlcNAcylation impairs endothelial cell functions

Senescent cells are characterized by a reduced capacity to proliferate in response to growth factors due to cell cycle arrest. Consistent with this, we observed a substantial reduction of bFGF-induced BrdU incorporation in endothelial cells with GFAT1/OGT double knockdown (Fig. 3A). Furthermore, the angiogenic capacity was decreased in O-GlcNAc-depleted cells. This was demonstrated in spheroid assays using VEGF as an angiogenic stimulus (Fig. 3B, C). In these assays, the number of capillary-like structures (sprouts) per spheroid induced by VEGF was reduced by 33% in cells with GFAT1/OGT double knockdown compared to control cells (Fig. 3D). Furthermore, VEGF-induced sprout length was reduced by ∼20% in O-GlcNAc-depleted cells (Fig. 3B, E). Overall, combining these two features resulted in a 37.4% reduction of the sum of sprout lengths per spheroid in response to VEGF (Figure 3B, F). As with the genetic approach, the use of the pharmacological agent OSMI-1 to deplete O-GlcNAc significantly decreased angiogenesis, leading to a 62.4% reduction in VEGF-induced sprouting compared to control cells (Fig. 3G, H). The reduced proliferative and angiogenic capacities observed here are consistent with the higher proportion of senescent cells in O-GlcNAc-depleted cultures and demonstrate the functional relevance of senescence triggered by O-GlcNAc depletion.

**Figure 3.**
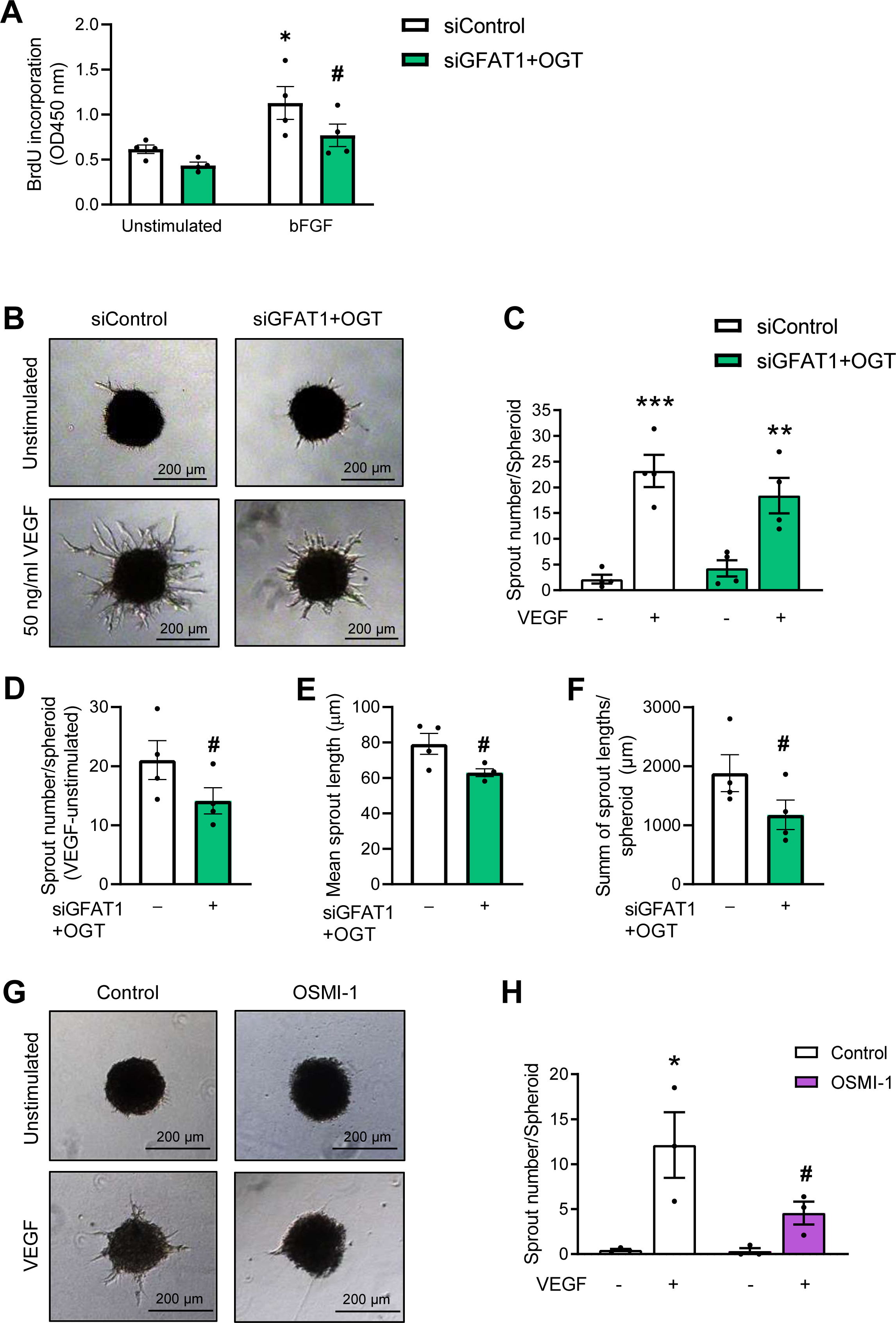
Reduced O-GlcNAcylation impairs proliferative and angiogenic function of endothelial cells. **(A)** HUVEC were repeatedly transfected with non-targeting siRNA (siControl) or co-transfected with GFAT1- and OGT-specific (siGFAT1+OGT) siRNAs (96h, twice) and seeded in full growth medium for BrdU assay; after 24 h, medium was changed for low serum (2% FCS) medium and after additional 4 h, cells were either treated with bFGF (50 ng/ml, 24h) or left unstimulated. **(B, G)** Spheroids generated from HUVEC depleted for O-GlcNAc using either genetic **(B)** or pharmacological **(G)** approach were treated with 10 or 50 ng/ml VEGF, respectively, or left unstimulated. Representative images of spheroids **(B, G)** and analysis of the number of sprouts per spheroid (C, D, H), mean sprout lengths **(E)** and sum of sprout lengths **(F)** are shown. Data are means ± SEM of 4 independent experiments using cells from different donors. *p < 0.05, **p < 0.01, ***p < 0.001 vs. respective unstimulated control (-); ^#^p < 0.05 vs. control siRNA or diluent control (**H**). Two-way ANOVA with Bonferroni’s multiple comparison test **(A, C, H)**, or paired t-test **(D-F)** were used.

### Mass spectrometry-based map of O-GlcNAcylated proteins in endothelial cells

To elucidate the mechanisms linking reduced O-GlcNAcylation to the induction of endothelial senescence, we next sought to identify O-GlcNAc-modified proteins in endothelial cells. To this end, we applied two complementary strategies of O-GlcNAc immunoenrichment.

To obtain a global view of the HUVEC O-GlcNAcome, we used a recently developed and commercially available mix of monoclonal antibodies (54) to enrich O-GlcNAcylated peptides following the tryptic digestion of HUVEC cells that had been treated with the OGA inhibitor Thiamet G to increase abundance of O-GlcNAc on the proteins and facilitate immunoenrichment (Fig. 4A). Upon collision-induced fragmentation on a trapped ion mobility – mass spectrometer, we identified about 850 HexNAc-modified peptide spectrum matches (PSMs) per measurement and reproducibly detected over 300 HexNAc peptides in at least two of three replicates, mapping to 187 modified proteins (Fig. 4B, Suppl. Table 1). In line with our previous data on synthetic O-GlcNAcylated peptides (45), these were shifted towards higher m/z values and lower ion mobility values as compared to unmodified peptides (Suppl. Fig. 2A). This correlates with the increased mass through the addition of the HexNAc moiety, and we also found modified peptides to be longer than unmodified peptides in our dataset, despite a slightly lower miscleavage rate (Suppl. Fig. 2B, C). This observation aligns with the semi-consensus sequence for O-GlcNAc, which implies a reduced probability of lysine and arginine residues around O-GlcNAc sites (55). Overlaying endothelial HexNAc-modified proteins from our dataset with the SeneQuest database (56) revealed 32 senescence-associated proteins (Fig. 4C). Clustering them using the STRING network analysis positioned β-catenin (CTNNB1) centrally, interacting with further transcriptional regulators including FOXO1, EGR1 and MITF. This observation aligns with the Gene Ontology enrichment analysis of the entire dataset, which shows a significant overrepresentation of proteins involved in transcriptional regulation (Fig. 4D).

**Figure 4.**
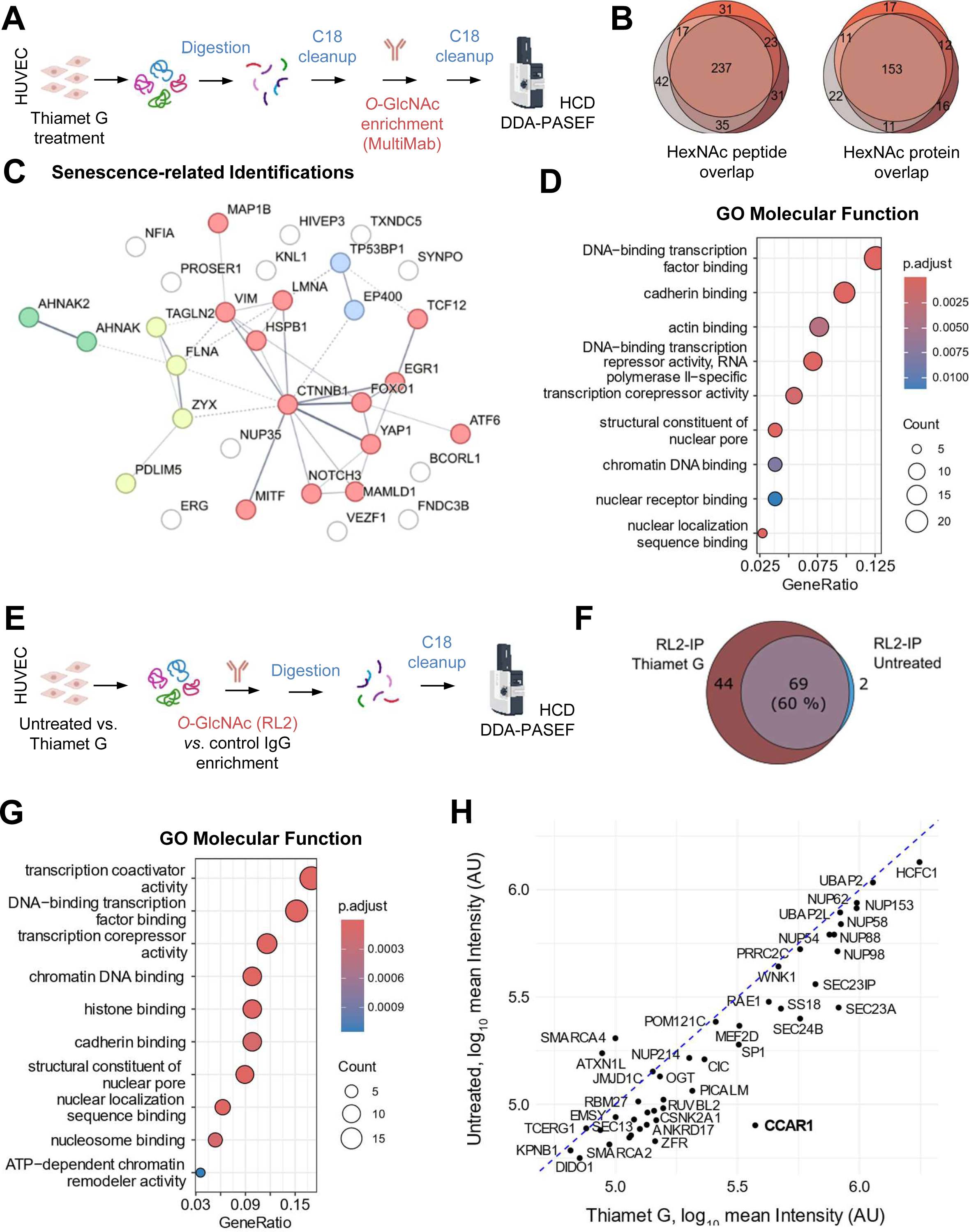
Identification of endothelial O-GlcNAc targets. **(A)** Human umbilical vein endothelial cell (HUVEC) lysates (Thiamet G-treated) were subjected to O-GlcNAc peptide-level enrichment using an antibody mix. **(B)** Overlap of unique HexNAc peptide and corresponding protein identifications from biological triplicates (Glycan FDR < 0.05). **(C)** HexNAc-modified proteins detected in at least two of three replicates (N = 187), overlapping with senescence-associated proteins from the SeneQuest database (filtered for proteins up- or downregulated in ≥3 studies, N = 893). The protein-protein interaction network was visualized using STRING with MCL clustering. **(D)** Gene Ontology (GO) enrichment for Molecular Function of HexNAc-modified proteins identified in at least two of three replicates (N = 187), shown in a simplified form. **(E)** Lysates from basal or O-GlcNAc-enriched HUVEC (Thiamet G-treated vs. untreated) were immunoprecipitated using the RL2 antibody or a control IgG and analyzed by LC-MS. **(F)** Venn diagram showing proteins identified in all three biological replicates, excluding identifications from the IgG control IP. **(G)** Gene Ontology (GO) enrichment for Molecular Function of HexNAc-modified proteins identified in at least two of three replicates (N = 187), shown in a simplified form. **(H)** Mean intensities (max-LFQ, n=3) of immunoprecipitated proteins comparing Thiamet G treatment vs. vehicle control (untreated). Only proteins quantified in all three replicates in both groups are shown.

Our second enrichment strategy was performed at the protein level to identify O-GlcNAcylated proteins with the well-characterized RL2 antibody (57, 58). Specifically, we used Thiamet G-treated vs. untreated HUVEC to identify substrates sensitive to O-GlcNAc modulation (Fig. 4E). Across three biological replicates, we identified about 1000 proteins of which we only retained those detected in all replicates and not in the nonspecific IgG control IP (Fig. 4F). This stringent filtering yielded 115 proteins, 38 % of which appeared only after Thiamet G treatment, while 60 % were detected in both conditions (Fig. 4F, Suppl. Table 2). GO-analysis revealed an enrichment pattern similar to the one observed with the peptide-based approach, dominated by terms related to transcriptional regulation and the nuclear pore complex (Fig. 4G). This finding is consistent with the origin of the RL2 antibody, which was raised against a nuclear pore complex protein fraction (58). Despite enriching O-GlcNAc at the protein level, we identified 112 unique HexNAc peptides (FragPipe glycan q-value <0.01): 61 in untreated and 102 in Thiamet G-treated cells, corresponding to 33 proteins (∼30% of the identified candidates). To visualize abundant targets and their sensitivity to Thiamet G, we compared the mean protein intensities from both conditions pairwise (Figure 4H). The mean protein intensities fold-change was approximately 1.5, with the largest changes upon Thiamet G-treatment observed for SEC24C (9.2-fold) and CCAR1 (4.7-fold), pointing at their high sensitivity to O-GlcNAc modulation.

### CCAR1 is a bona fide O-GlcNAc substrate in endothelial cells

Given the high sensitivity of CCAR1 to O-GlcNAc modulation together with its reported roles in senescence-related processes, such as the cell cycle (59), we decided to investigate its O-GlcNAcylation and functional relevance in human endothelial cells further. We immunoprecipitated CCAR1 from either untreated or Thiamet G-treated HUVEC and analyzed the immunoprecipitates by Western blot using an O-GlcNAc-specific antibody (CTD110.6). A basal O-GlcNAc signal was detected at the expected molecular weight of CCAR1, which increased markedly upon Thiamet G treatment (Fig. 5A). This finding confirms the profound enrichment of CCAR1 observed in our RL2-based O-GlcNAc screen after Thiamet G treatment. However, the sequence coverage of CCAR1 in our MS analysis was only 50%, which we attributed, at least in part, to the low frequency of arginine and lysine residues in the N-terminal domain. To improve sequence coverage, we digested CCAR1 immunoprecipitates with chymotrypsin, which resulted in the identification of eight HexNAc-modified peptides (Fig. 5B). The two most abundant modified peptides “TATAVSQPAALGVQQPSLL[HexNAc]” and “TQPAVALPTSL[HexNAc]” were reproducibly detected in all six samples and indicated an increased occupancy upon OGA inhibition (1.5-fold and 2-fold, respectively) (Fig. 5C-E).

**Figure 5.**
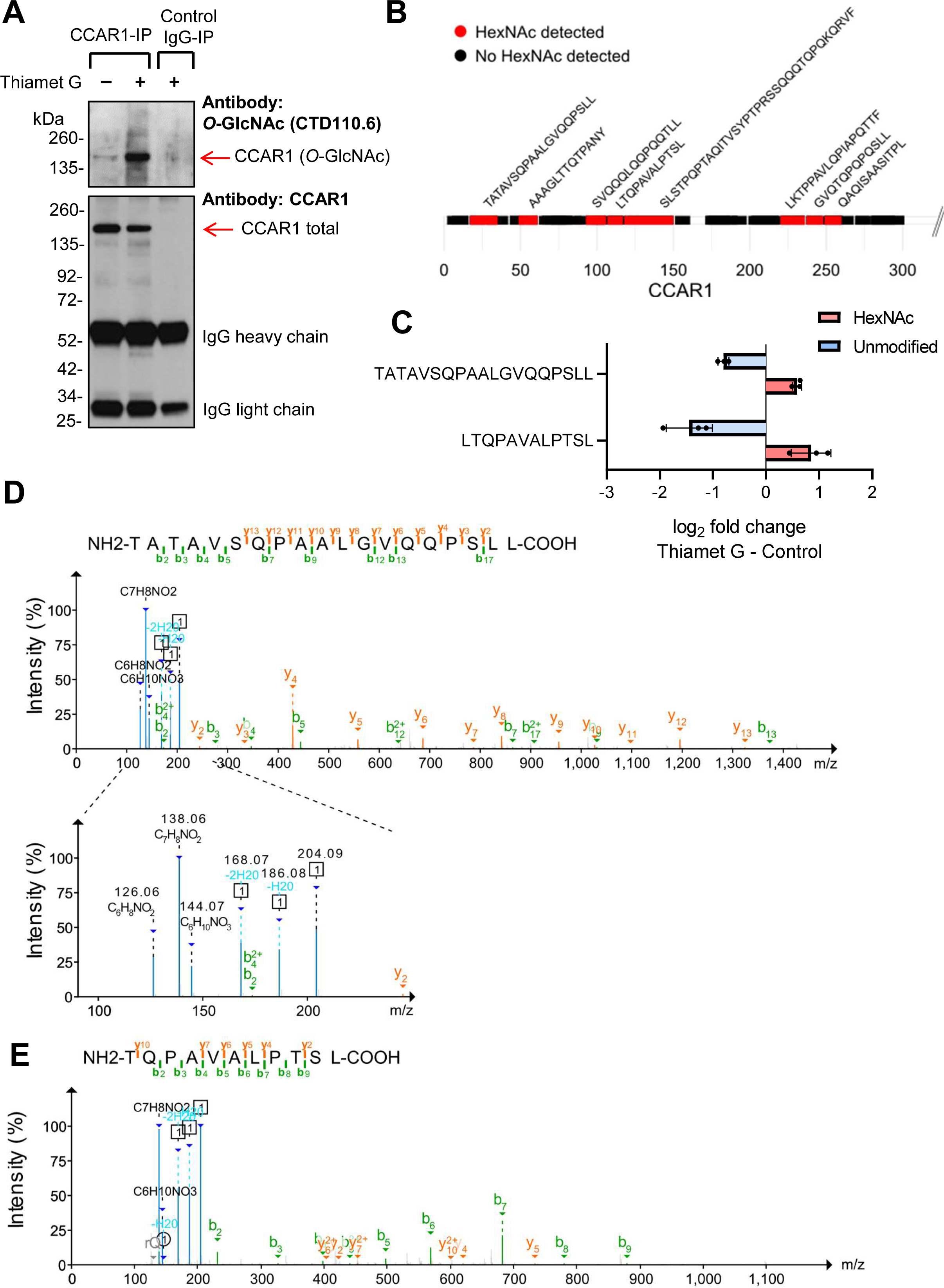
Identification and validation of CCAR1 O-GlcNAcylation. **(A)** Lysates of control or Thiamet G-treated HUVEC were immunoprecipitated for CCAR1 and stained by immunoblot for O-GlcNAc (upper panel) and CCAR1 (lower panel). Images are representative of 3 independent experiments using cells from different donors. **(B)** MS-based analysis of enriched proteins after CCAR1 co-immunoprecipitation. Coverage of the N-terminal region of CCAR1 by LC-MS using chymotrypsin as protease. HexNAc modified peptides are indicated in red. **(C)** MS-intensity fold change between Thiamet G-treatment and control, normalized to protein fold change, for the two most abundant HexNAc peptides and their unmodified counterpart. Data are means ± SD of 3 independent experiments using cells from different donors. **(D, E)** Representative MS/MS spectrum of the HexNAc modified peptide “TATAVSQPAALGVQQPSLL” **(D)** and “TQPAVALPTSL” **(E)**. Peak colors: orange, green and blue refer to y, b and diagnostic oxonium ions, respectively. The enlarged view in the m/z range of 100 to 225 **(D)** shows the [HexNAc]+ ion (C8H14NO5) and corresponding fragmentation induced oxonium ions.

### CCAR1 is involved in apoptosis and DNA damage regulation in endothelial cells

CCAR1 is a poorly characterized protein and its function in endothelial cells remains unknown. CCAR1 has been reported to regulate the cell cycle, apoptosis and the DNA damage response (59), processes chiefly involved in senescence. In order to investigate whether CCAR1 also regulates these functions in endothelial cells, we transfected HUVEC with CCAR1-specific siRNA in two consecutive 96-h transfection rounds. This approach resulted in the efficient downregulation of CCAR1 by 71% (Figure 6A). However, under these conditions, we did not observe any significant change in canonical markers of senescence signaling (Fig. 6A), which prompted us to examine the key components of apoptotic signaling. We found that CCAR1 depletion resulted in considerable downregulation of caspase 3, a central effector caspase in apoptosis, and almost completely prevented staurosporine-induced caspase 3 activation (Fig. 6B). With respect to DNA damage, CCAR1 knockdown alone did not activate the DNA damage response signaling. However, it significantly sensitized cells to H_2_O_2_-induced DNA damage, as demonstrated by increased phosphorylation of p53, Chk-1 and H2A.X (Fig. 6C). Thus, while CCAR1 depletion alone may be insufficient to induce senescence, it could exacerbate endothelial senescence under stress conditions, while simultaneously increasing resistance to apoptosis, another hallmark of senescence. Based on these observations, we concluded that CCAR1 may protect cells from stress-induced senescence.

**Figure 6.**
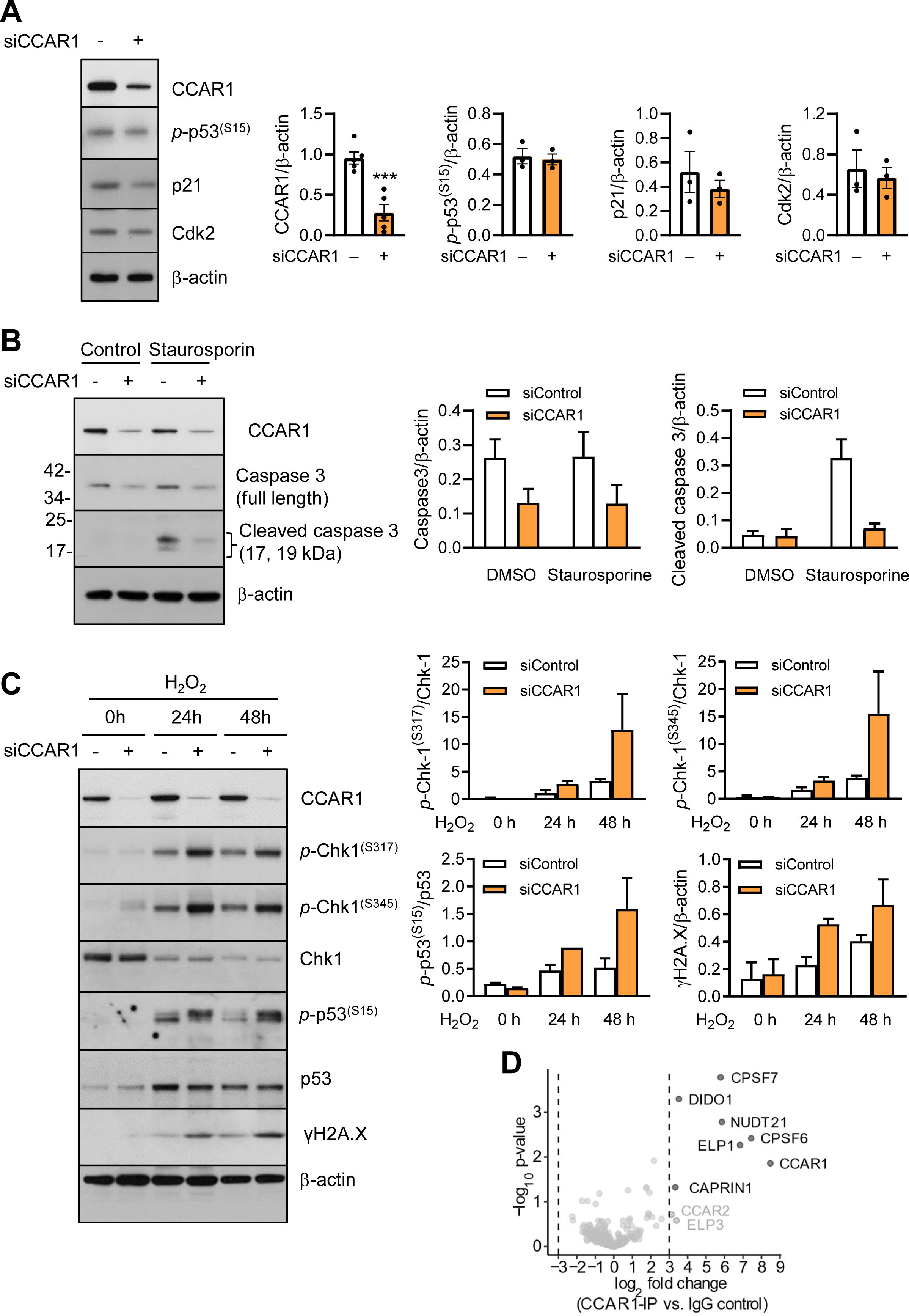
CCAR1 knockdown interferes with apoptotic and DDR signaling in endothelial cells. **(A, B)** HUVEC were repeatedly transfected with non-targeting siRNA (-) or transfected with CCAR1-specific siRNA (siCCAR1, +) (96h, twice). Afterwards cells were harvested for immunoblot analysis **(A)** or treated either with staurosporin (2 µM, 2h) or vehicles (control) followed by harvesting for immunoblot **(B). (C)** 48h post 2^nd^ transfection round, cells were treated with H_2_O_2_ for the indicated periods of time and harvested for immunoblot analysis. Representative immunoblots and densitometrical analyses are shown. Data are means ± SEM of 2 independent experiments using cells from different donors. **(D)** Identification of potential interaction partners after CCAR1-immunoprecipitation and trypsin digestion. Volcano plot of the intensity fold change (CCAR1-IP vs. control IgG-IP). Three independent experiments using cells from different donors were performed. Data were filtered to contain at least 3 valid values in one of the groups. Missing values were imputed from the normal distribution.

To find a mechanistic explanation for these functional outcomes, we analyzed CCAR1 co-immunoprecipitates by MS (Fig. 6D). Seven strong candidates for CCAR1 interaction partners were identified, including the well-known CCAR1 interactor cell cycle-associated protein 1 (CAPRIN1). Among the novel potential CCAR1interaction partners, we identified DIDO1, ELP1 and components of the cleavage factor Im complex (CPSF6, CPSF7, NUDT21). All of these proteins have previously been implicated in apoptosis and DNA regulation, which is consistent with the role of endothelial CCAR1 in these processes, as demonstrated here.

### CCAR1 overexpression attenuates premature stress-induced endothelial senescence

To gain a better understanding of whether CCAR1 plays a protective role in stress-induced senescence, we investigated CCAR expression and O-GlcNAcylation in our premature senescence model. We observed a significant reduction in total CCAR1 levels in cells, in which senescence was induced by exposure to oxidative stress (Fig. 7A). Furthermore, CCAR1 immunoprecipitated from prematurely senescent cells exhibited reduced O-GlcNAcylation compared to controls when the immunoprecipitates were loaded in amounts normalized for total CCAR1 levels (Figure 7B). These data suggest that O-GlcNAcylation may regulate the stability of CCAR1.

**Figure 7.**
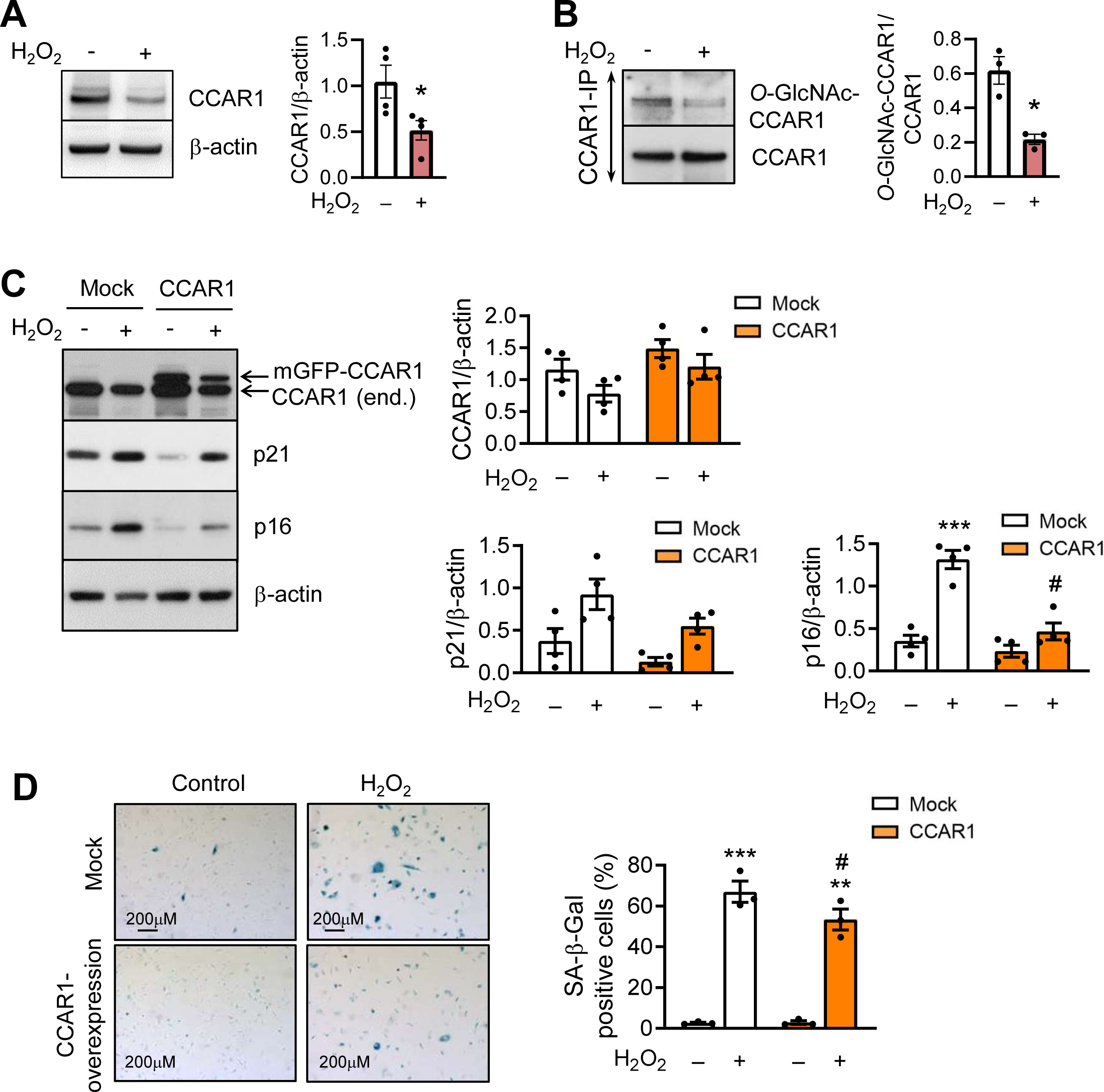
CCAR1 overexpression ameliorates stress-induced senescence. **(A, B)** Cells were treated either with H_2_O_2_ (100 µM, 8 days) or vehicle as control (-). Afterwards, cells were harvested for immunoblot analysis **(A)** or for CCAR1 immunoprecipitation **(B)**. **(C D)** Untransduced control cells (mock) or CCAR1-overexpessing HUVEC obtained from the same donor were treated with H_2_O_2_ as described above to induce premature senescence. Cells were analyzed using SA-β-gal staining or harvested for immunoblotting. Representative immunoblots **(A-C)** and light microscopy images **(D)** with corresponding analyses are shown. Data are means ± SEM of 3–4 independent experiments using cells from different donors. *p<0.05 vs. control, paired t-test **(A, B)**. ***p<0.001 vs. control, ^#^p<0.05 vs. respectively treated mock, two-way ANOVA with Bonferroni’s multiple comparison test **(C, D)**.

We then generated HUVEC that stably expressed mGFP-CCAR1 within a physiological range, which at this culture stage resulted in a 1.3-fold increase in CCAR1 levels in transductants compared to untransduced control cells under basal conditions and which was sufficient to keep CCAR1 levels replenished in transductants upon H_2_O_2_ treatment (Fig. 7C). When transduced cells were treated with H_2_O_2_ to induce premature senescence, reduced levels of p21 and p16, as well as decreased SA-β-gal staining was observed (Figure 7C, D) when compared to untransduced controls. Thus, restoring CCAR1 levels in H_2_O_2_-treated cells partially attenuated senescence supporting a protective role for CCAR1 in premature oxidative stress-induced senescence.

### CCAR1 overexpression protects endothelial cells from replicative senescence

To investigate whether CCAR1 also protects against chronological aging, we assessed its levels in replicative senescent HUVEC. We observed no difference in CCAR1 levels up to P20 (Fig. 8A), which is the condition that we used for the replicative senescence experiments described above. However, extending the experiments to later passages revealed reduced CCAR1 levels at passage 25 (P25) and strongly reduced CCAR1 levels at passage 30 (P30) (Fig. 8B, C). These results suggest that the CCAR1 depletion may predominantly affect later stages of senescence, possibly by stabilizing the senescence phenotype.

**Figure 8.**
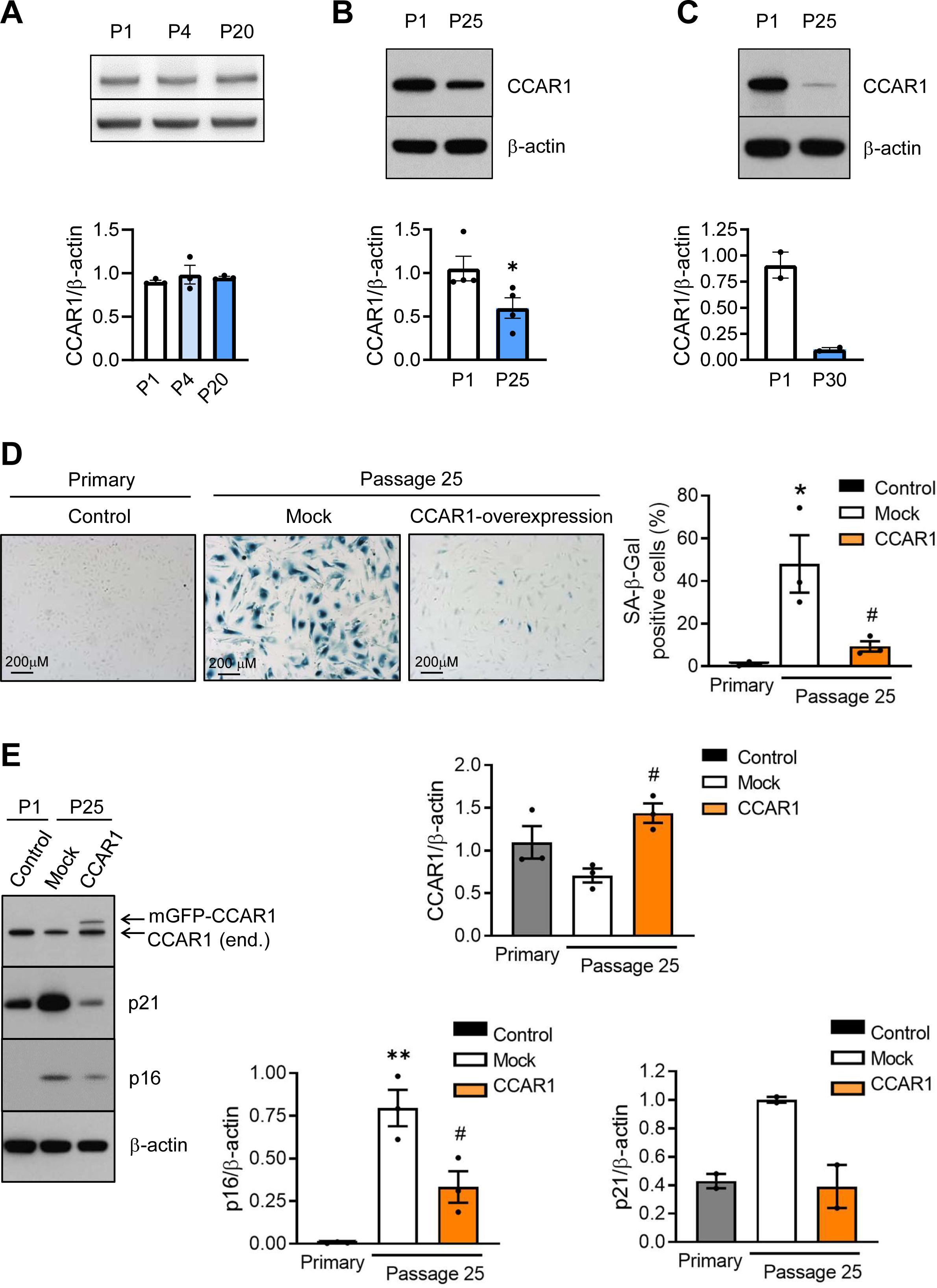
CCAR1 acts as a protective player at the later stages of replicative senescence. **(A-C)** CCAR1 levels are significantly decreased at later senescence stages. **(D-E)** Untransduced control cells (mock) or CCAR1-overexpessing HUVEC obtained from the same donor were continuously cultured up to passage 25 and finally analyzed using SA-β-gal staining **(D)** or immunoblot **(E)**, P1 HUVEC, representing a separate biological replicate, were processed in parallel. Representative immunoblots **(A-C, E)** and light microscopy images **(D)** with corresponding analyses are shown. Data are means ± SEM of 3 independent experiments using endothelial cells from different donors. *p<0.05 vs. P1 control, unpaired t-test **(B, C)**. *p<0.05, **p<0.05 vs. P1 control, #p<0.05 vs. untransduced, one-way ANOVA with Bonferroni’s multiple comparison test **(D, E)**.

To test whether CCAR1 overexpression attenuates the onset of replicative senescence, we subjected non-transduced controls and HUVEC overexpressing CCAR1 to repeated passaging. Strikingly, CCAR1 overexpression (approximately twofold at P25) was found to significantly protect cells from replicative senescence. At P25, control cells exhibited a pronounced senescent phenotype, as evidenced by increased β-gal staining (Fig. 8D) and the induction of the biochemical senescence markers p21 and p16 (Fig. 8E). By contrast, CCAR1-overexpressing cells generated from the same donor as control cells maintained a morphology characteristic of young cells and the levels of senescence markers remained comparable to those observed in P1 cells (Fig. 8D, E). To our knowledge, these data provide the first evidence of a protective role for CCAR1 in endothelial senescence.

## Discussion

Altered O-GlcNAcylation is increasingly recognized as a hallmark of age-related pathologies (9, 60). These include type 2 diabetes and its vascular complications, where hyperglycemia-induced endothelial dysfunction, driven by elevated O-GlcNAc levels, plays a mechanistic role (61). Aging, a major risk factor for vasculopathies, is itself associated with dysregulated O-GlcNAcylation (9). However, the mechanisms linking O-GlcNAc dysregulation to vascular aging and related abnormalities, as well as the temporal sequence of these interconnected events, remain unclear. Elucidating these mechanisms is crucial, as maintaining O-GlcNAc homeostasis could provide new therapeutic approaches for preventing or treating age-related diseases (38). In this study, we show that inhibition of O-GlcNAcylation triggers senescence. We characterize the O-GlcNAcome of endothelial cells, and introduce CCAR1 as an O-GlcNAcylated substrates that plays a role in the regulation of endothelial senescence. By linking altered O-GlcNAcylation to specific substrates and components of senescence signalling, we establish an initial molecular connection between O-GlcNAc dysregulation, endothelial senescence and vascular aging.

Our first observation was that global O-GlcNAc levels decreased during both premature and replicative senescence in endothelial cells. This reduction was accompanied by decreased levels of GFAT, the rate-limiting enzyme of the HBP, in both models, indicating the suppression of HBP flux during endothelial senescence. In replicative senescence, additional changes in OGT and OGA levels suggest that altered O-GlcNAc cycling is another contributing factor to reduced O-GlcNAc levels. Of note, these alterations were evident starting from passage 4 onwards, suggesting that they may contribute to the development of the overt senescence phenotype. Overall, our data show that altered regulation of the O-GlcNAc system, resulting in decreased O-GlcNAcylation, is a characteristic feature of endothelial senescence.

In addition to changes in enzyme abundance, post-translational and transcriptional mechanisms may modulate O-GlcNAc levels during endothelial senescence. For example, AMPK-mediated phosphorylation of GFAT at serine 243 (62) could suppress HBP flux, particularly under conditions of oxidative stress, where AMPK activation is well documented (63–65). Furthermore, the transcriptional regulators of GFAT1, SP1 and XBP1 (66, 67) are known to decline with senescence and aging (68–70), which could contribute to GFAT downregulation.

In terms of the metabolic regulation of the HBP, a reduced availability of fructose-6-phosphate, the substrate derived from glycolysis, is unlikely, at least in replicatively senescent cells, which have been previously shown to exhibit increased glycolytic activity (71). However, channeling glutamine towards energy production due to increased glutaminolysis, as reported in replicatively senescent HUVEC (72), could limit its availability for GFAT and thereby constrain HBP flux, despite sufficient glycolytic input.

Our next data provided evidence that reduced O-GlcNAcylation is a driving force, rather than a correlative marker, of endothelial senescence. We found that reducing O-GlcNAcylation through genetic or pharmacological approaches mimics the molecular characteristics and functional impairments of endothelial senescence, including diminished proliferation and reduced angiogenic capacity. Interestingly, this senescence phenotype developed independently of the canonical DNA damage response involving the phosphorylation of H2A.X (serine 139) or p53 (serine 15) (73, 74). Instead, elevated p53 phosphorylation at serine 46 accompanied by ERK activation suggests the engagement of an alternative MEK-ERK-dependent pathway that reinforces the senescence program through p53 and the p16-pRb axis (75–78).

As elevated O-GlcNAcylation has been linked to glucose toxicity and vascular dysfunction under hyperglycemic conditions (79–82), we initially anticipated that lowering O-GlcNAc levels would attenuate senescence. Instead, our findings revealed that, under physiological glucose conditions, maintenance of O-GlcNAc homeostasis is essential for endothelial resilience, and its disruption may destabilize protective programs and favor senescence. By delivering the first comprehensive map of the human endothelial O-GlcNAcome, we show that O-GlcNAcylation contributes to endothelial homeostasis by targeting components of gene-expression networks. Consistent with the meta-analysis by Wolff-Fuentes et al. (55), this highlights the dominant role of O-GlcNAc in genomic regulation. Extending the view of this network-level regulation, our data reveal that O-GlcNAcylation directly targets multiple factors involved in chromatin remodeling and methylation control – processes that are closely associated with senescence and aging. Our analysis identified 187 O-GlcNAc-modified proteins, many of which have also been reported in other cell types, though their functional significance remains largely unexplored. Here, we provide experimental evidence for their O-GlcNAcylation in human endothelial cells. For example, we detected O-GlycNAcylation of lysine-specific demethylase 3B (KDM3B), PROSER1, and the transcription factor VEZF1, which influence histone and DNA methylation dynamics (83–86). We also identified transcriptional and post-transcriptional regulators such as MITF, which activates p16 and promotes retinoblastoma hypophosphorylation (87), and RBMS2, which stabilizes p21 mRNA (88). O-GlcNAcylation of such factors may link O-GlcNAc signaling to the activation of senescence pathways beyond the canonical DNA-damage response as suggested from our data. In addition to nuclear and epigenetic regulators, several of the most abundantly O-GlcNAc-modified proteins point to broader effects of O-GlcNAc signaling on cell-cycle control and proliferation. Among these, we identified PRRC2C (89), MGA (90), RAPH1 (91), and the RNA-binding protein ZFR (92), whose hyper-O-GlcNAcylation was shown to promote endothelial proliferation under hyperglycemia (92). As expected, multiple nucleoporins, which are classical O-GlcNAc substrates, were also highly modified. Their emerging role in genome regulation and involvement in age-related diseases (93) adds functional relevance to these findings. Finally, detection of EGF domain-specific OGT (EOGT) substrates NOTCH2, NOTCH3, and STAB1 indicates that cytosolic O-GlcNAc metabolism can indirectly influence extracellular pathways maintaining vascular integrity, as EOGT in the endoplasmic reticulum depends on cytosolic UDP-GlcNAc (94, 95).

Interestingly, none of the well-established “classical” endothelial O-GlcNAc targets was detected in our screen, including eNOS (13, 14, 96), SP1 (97), Akt (98), or components of insulin signaling such as IRS-1/2 and PI3K (14). These proteins are typically hyper-O-GlcNAcylated under hyperglycemic conditions, which promotes vascular dysfunction and insulin resistance (reviewed in (61, 99)). Therefore, we speculate that the endothelial O-GlcNAcome detected under physiological glucose conditions and described here may represent a steady-state regulatory network of O-GlcNAc signaling, distinct from the stress-induced modifications prevalent in hyperglycemic or diabetic states.

In this context, individual O-GlcNAc-modified proteins may act as key effectors that translate metabolic cues into senescence-associated signaling. CCAR1 (cell division cycle and apoptosis regulator 1) is one such protein that has emerged as a particularly compelling candidate. Its robust O-GlcNAcylation under Thiamet G treatment confirmed for the first time its status as a bona fide endothelial substrate. The identified modification sites cluster within the N-terminal region of CCAR1 (comprising ∼260 amino acids), which contains an S1-like RNA-binding domain (100, 101) and a predicted nuclear localization sequence (100). This suggests that O-GlcNAcylation within this segment may modulate transcriptional and nuclear regulatory activities of CCAR1.

Although CCAR1 was originally characterized for its role in cell-cycle regulation, transcriptional control, and apoptosis (102–104), its function remains incompletely understood. Here, we reveal an unrecognized protective function of CCAR1 in endothelial senescence: its expression declines during both oxidative stress-induced and replicative senescence, and restoring it mitigates premature senescence and delays replicative aging. These protective effects appear to result from CCAR1’s role in preserving genomic stability and coordinating stress-response pathways. We show that depletion of CCAR1 sensitized endothelial cells to oxidative stress, as evidenced by enhanced phosphorylation of DNA damage response mediators such as p53, Chk-1, and H2A.X. Since CCAR1 levels decline with replicative age, this sensitization may amplify cumulative DNA damage over successive divisions, promoting irreversible cell-cycle arrest. This observation is consistent with a recent report that CCAR1 contributes to genome maintenance by regulating FANCA mRNA splicing to sustain DNA repair capacity (105). Consequently, its loss may therefore compromise repair mechanisms and accelerate senescence under sustained stress.

Another mechanism by which CCAR1 may influence endothelial senescence is its role in regulating apoptosis. It has been demonstrated that CCAR1 depletion confers resistance to cytotoxic and DNA-damaging agents (102, 106). Consistently, our study found that CCAR1 knockdown in HUVEC attenuated staurosporine-induced caspase 3 activation. As resistance to apoptosis is a hallmark of senescence (107), downregulation of CCAR1 in senescent endothelial cells may confer a survival advantage. Supporting this view, our proteomic analysis identified CCAR1-associated complexes involved in apoptosis, RNA processing, and cell-cycle regulation. Taken together, these findings suggest that CCAR1 acts as a nodal integrator that couples the programmes of DNA repair, apoptosis, and senescence to maintain endothelial homeostasis and to delay endothelial aging. However, determining the functional role of the individual O-GlcNAc modification sites of CCAR1, particularly in the context of endothelial senescence, will be an important focus of future research.

In summary, we present a draft of the human vascular endothelial O-GlcNAcome, newly associating substrates such as CCAR1 with senescence pathways. Our study not only demonstrates that reduced O-GlcNAcylation is mechanistically linked to endothelial senescence and associated functional impairments, but also uncovers a network of constitutively O-GlcNAcylated proteins that sustain endothelial homeostasis. These findings establish O-GlcNAc signaling as a promising target for counteracting endothelial dysfunction and maintaining vascular health during aging.

## Authors’ contributions

FM and DZ conceived the study. RH, FM and DZ supervised the work. AW and DZ designed and performed the experiment, and analyzed the data. CE and FRS provided valuable materials and helpful advices. AW, RH, FM and DZ wrote the manuscript. All authors discussed the results and commented on the manuscript.

## Acknowledgements

We are grateful to Elke Teuscher (Institute for Molecular Cell Biology, Center for Molecular Biomedicine, Jena University Hospital) for excellent assistance with the isolation and culture of HUVEC.

---

**Supplementary Figure S1.**
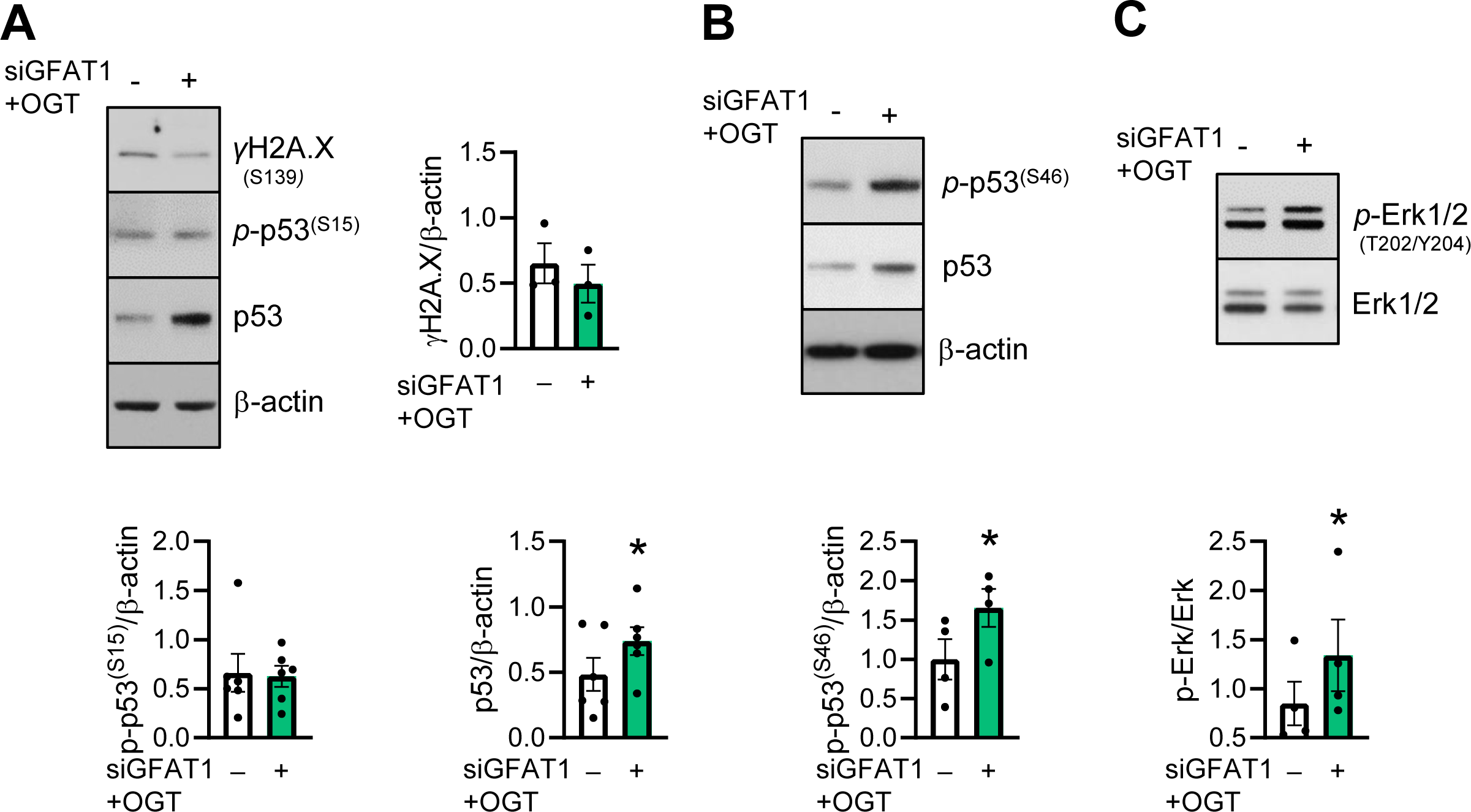
O-GlcNAc depletion stabilizes p53 via serine 46 phosphorylation. **(A-C)** HUVEC were repeatedly transfected with non-targeting siRNA (-) or co-transfected with GFAT1- and OGT-specific siRNAs (siGFAT1+OGT) (96h, twice). Representative immunoblots and densitometric analyses are shown. Data are means ± SEM of 4-6 independent experiments using cells from different donors. *p<0.05, vs. control, paired t-test.

**Supplementary Figure S2.**
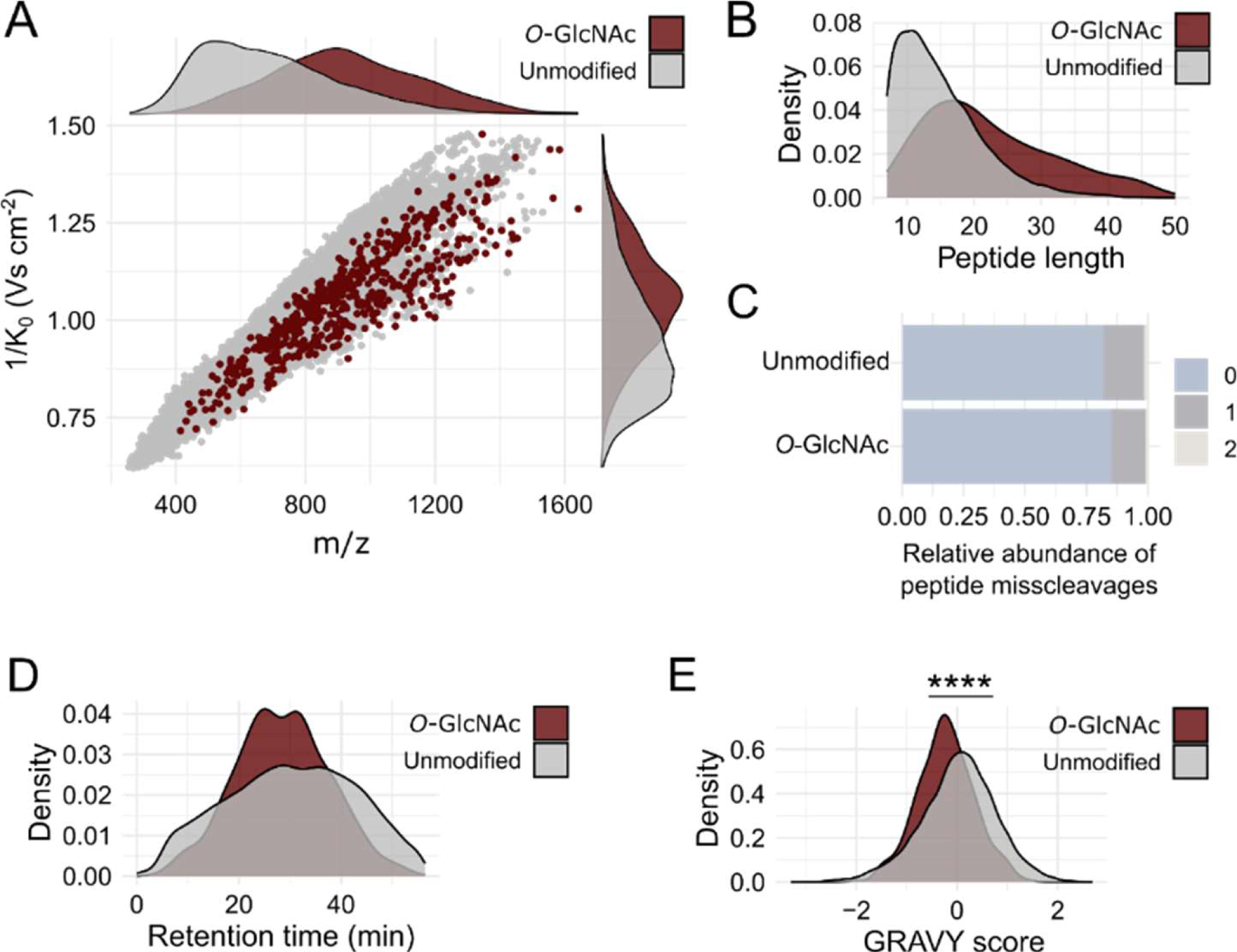
O-GlcNAcylated peptides are shifted in the m/z vs. ion mobility dimension. O-GlcNAc-enriched tryptic peptides (N=3) were measured by LC-TIMS/MS with HCD fragmentation. The analysis was performed on the PSM level. **(A)** Distribution of unmodified and HexNAc-modified PSMs in the ion mobility (1/K0) vs. m/z dimension. Shown are unique peptide sequences for each charge state (Unmodified: N = 23,157 HexNAc: N = 505). **(B)** Relative abundance of peptide misscleavages for unmodified and HexNAc-modified PSMs. **(C-E)** Unmodified and HexNAc-modified PSMs were compared. Shown are unique peptide sequences, with the highest observed intensity (Unmodified: N = 18,73867; O-GlcNAc: N = 416). **(C)** Peptide length distribution. **(D)** Retention time distribution. **(E)** Distribution of the GRAVY score as a metric of hydrophobicity, with a higher score indicating higher hydrophobicity. Asymptotic two-sample Kolmogorov-Smirnov-test, **** p<1e-04.

