## Supplementary material for "Mapping the endothelial O-GlcNAcome uncovers CCAR1 as a regulator of senescence": Suppl Table 1

| Peptide | Gene | Protein | _log2_Int | ean_Intensi | CV | _datase |
|---|---|---|---|---|---|---|
| QQSSSLTSVPPTTF! | PRRC2C | sp\|Q9Y520\|PRC2C_HUMAN | 25,11 | 3,64E+07 | 7,55 | 3 |
| QLFVTVVK | ANKRD17 | sp\|O75179\|ANR17_HUMAN | 23,56 | 1,25E+07 | 11,03 | 3 |
| ISPPEPQSFASK | MGA | sp\|Q8IWI9\|MGAP_HUMAN | 23,45 | 1,14E+07 | 5,41 | 3 |
| \QSALSPYLSSPVSS | RBMS2 | sp\|Q15434\|RBMS2_HUMAN | 23,44 | 1,15E+07 | 15,32 | 3 |
| VPSSFMLPIR | HIVEP1 | sp\|P15822\|ZEP1_HUMAN | 23,05 | 8,71E+06 | 1,42 | 3 |
| PLVTIPAPTSTK | RAPH1 | sp\|Q70E73\|RAPH1_HUMAN | 22,98 | 8,44E+06 | 24,13 | 3 |
| ATQSNEITIPVTFES | HSPB1 | sp\|P04792\|HSPB1_HUMAN | 22,96 | 8,18E+06 | 12,81 | 3 |
| RQLFVTVVK | ANKRD17 | sp\|O75179\|ANR17_HUMAN | 22,86 | 7,84E+06 | 27,06 | 3 |
| VVSTLPSTVLGK | GMEB2 | sp\|Q9UKD1\|GMEB2_HUMAN | 22,59 | 6,32E+06 | 4,04 | 3 |
| VSTTAPVTLASSK | JMJD1C | sp\|Q15652\|JHD2C_HUMAN | 22,31 | 5,24E+06 | 12,25 | 3 |
| CQPVFSFGNSEQTI | NUP153 | sp\|P49790\|NU153_HUMAN | 22,29 | 5,15E+06 | 12,72 | 3 |
| MIPICPVVSFTYVPS | ZFR | sp\|Q96KR1\|ZFR_HUMAN | 22,21 | 4,84E+06 | 3,02 | 3 |
| PTTVVPLISTIAGDS | VEZF1 | sp\|Q14119\|VEZF1_HUMAN | 21,96 | 4,09E+06 | 8,04 | 3 |
| LGGSGSCFCDEGW | STAB1 | sp\|Q9NY15\|STAB1_HUMAN | 21,80 | 3,68E+06 | 15,32 | 3 |
| SSFASQASGSSSSA | KDM3B | sp\|Q7LBC6\|KDM3B_HUMAN | 21,51 | 3,00E+06 | 11,48 | 3 |
| TLHFPTSPIIQQPG | NFIA | tr\|S4R3W2\|S4R3W2_HUMAN | 21,43 | 2,87E+06 | 19,90 | 3 |
| PPPPPPIPAPLPPQ | RAPH1 | sp\|Q70E73\|RAPH1_HUMAN | 21,39 | 2,78E+06 | 19,70 | 3 |
| TSDLHISSTPAATT | PROSER1 | sp\|Q86XN7\|PRSR1_HUMAN | 21,28 | 2,57E+06 | 15,35 | 3 |
| AINVSVPTTLPSATC | MITF | sp\|O75030\|MITF_HUMAN | 21,28 | 2,59E+06 | 24,49 | 3 |
| YSCVCSPGFTGQR | NOTCH2 | sp\|Q04721\|NOTC2_HUMAN | 21,22 | 2,48E+06 | 19,99 | 3 |
| FGGTGAPTGGFTF | NUP62 | sp\|P37198\|NUP62_HUMAN | 21,09 | 2,23E+06 | 10,37 | 3 |
| VPLSAYER | SRRM2 | sp\|Q9UQ35\|SRRM2_HUMAN | 20,98 | 2,07E+06 | 1,66 | 3 |
| SYVTTSTR | VIM | sp\|P08670\|VIME_HUMAN | 20,97 | 2,06E+06 | 5,88 | 3 |
| TTNYSTVPQK | ZC3H14 | sp\|Q6PJT7\|ZC3HE_HUMAN | 20,81 | 1,84E+06 | 6,45 | 3 |
| GSAPPTTYHPPLQ | YLPM1 | sp\|P49750\|YLPM1_HUMAN | 20,78 | 1,80E+06 | 6,09 | 3 |
| SLPASSYSQDPVYA | NOL4L | sp\|Q96MY1\|NOL4L_HUMAN | 20,53 | 1,52E+06 | 15,10 | 3 |
| APVTVTSLPAGVR | HCFC1 | sp\|P51610\|HCFC1_HUMAN | 20,51 | 1,52E+06 | 20,49 | 3 |
| SPITIITTK | HCFC1 | sp\|P51610\|HCFC1_HUMAN | 20,44 | 1,42E+06 | 4,49 | 3 |
| PPPPPATTQNYQD | ZFR | sp\|Q96KR1\|ZFR_HUMAN | 20,38 | 1,37E+06 | 18,04 | 3 |
| LFPSSLAGETLGSF | NUP214 | sp\|P35658\|NU214_HUMAN | 20,17 | 1,21E+06 | 25,98 | 3 |
| PPMDVMHSSVYC | RC3H2 | sp\|Q9HBD1\|RC3H2_HUMAN | 20,17 | 1,18E+06 | 3,77 | 3 |
| IAPTVTTWSNK | SEC31A | tr\|D6REX3\|D6REX3_HUMAN | 20,12 | 1,15E+06 | 15,34 | 3 |
| QKPTGTFSSGGGS\ | NUP214 | sp\|P35658\|NU214_HUMAN | 20,10 | 1,13E+06 | 18,76 | 3 |
| IVTTSPSSTFVPNIL | EMSY | sp\|Q7Z589\|EMSY_HUMAN | 20,08 | 1,11E+06 | 9,01 | 3 |
| YEANPGVPPATMS | SFPQ | sp\|P23246\|SFPQ_HUMAN | 20,05 | 1,10E+06 | 16,81 | 3 |
| CTPVPVTMTATPF | RPRD2 | sp\|Q5VT52\|RPRD2_HUMAN | 20,04 | 1,09E+06 | 21,94 | 3 |
| TYGATAETLSTSTT | EPB41L1 | \|A0A1B0GTW6\|A0A1B0GTW6_HUMA | 20,03 | 1,07E+06 | 3,89 | 3 |
| NTVSWEQSLFSTTI | KNL1 | sp\|Q8NG31\|KNL1_HUMAN | 20,01 | 1,14E+06 | 42,96 | 3 |
| MAVTPGTTTLPAT\ | HCFC1 | sp\|P51610\|HCFC1_HUMAN | 19,98 | 1,04E+06 | 11,69 | 3 |
| GLPTPPVPATLAYT | NBEAL2 | sp\|Q6ZNJ1\|NBEL2_HUMAN | 19,90 | 9,79E+05 | 5,62 | 3 |
| AGPSEPVTLASAG\ | BICRA | sp\|Q9NZM4\|BICRA_HUMAN | 19,85 | 1,00E+06 | 40,48 | 3 |
| SAIITQAGATGVTS | HCFC1 | sp\|P51610\|HCFC1_HUMAN | 19,83 | 9,32E+05 | 2,57 | 3 |
| ASTSDYQVISDR | NUP35 | sp\|Q8NFH5\|NUP35_HUMAN | 19,73 | 8,73E+05 | 9,21 | 3 |
| PLGTTLAVQAVPTA | ATF7IP | sp\|Q6VMQ6\|MCAF1_HUMAN | 19,70 | 8,54E+05 | 12,60 | 3 |
| VGSPATVTFQQNK | BPTF | sp\|Q12830\|BPTF_HUMAN | 19,66 | 8,29E+05 | 9,80 | 3 |
| GTGQPLPPLQIPQC | SEC31A | tr\|D6REX3\|D6REX3_HUMAN | 19,62 | 8,15E+05 | 16,76 | 3 |
| SISNPPGSNLRTT | WNK1 | sp\|Q9H4A3\|WNK1_HUMAN | 19,47 | 7,44E+05 | 26,42 | 3 |
| AFTPSSTMMEVFL( | UBAP2L | sp\|Q14157\|UBP2L_HUMAN | 19,39 | 6,85E+05 | 2,80 | 3 |
| LVNEAPVYSVYSK | STAM2 | sp\|O75886\|STAM2_HUMAN | 19,37 | 6,79E+05 | 6,65 | 3 |

| | | | | | | |
|---|---|---|---|---|---|---|
| TGTQATYTRPTVS | EMSY | sp\|Q7Z589\|EMSY_HUMAN | 19,29 | 6,43E+05 | 8,49 | 3 |
| SSGYVPPPVATPF | ZYX | sp\|Q15942\|ZYX_HUMAN | 19,27 | 6,53E+05 | 29,62 | 3 |
| STFEDVTQVSSAY | CTTN | sp\|Q14247\|SRC8_HUMAN | 19,25 | 6,25E+05 | 10,16 | 3 |
| TPPVTTNR | SBNO1 | sp\|A3KN83\|SBNO1_HUMAN | 19,24 | 6,23E+05 | 17,70 | 3 |
| DTLVNNVPLPNTL | PRRC2C | sp\|Q9Y520\|PRC2C_HUMAN | 19,23 | 6,21E+05 | 15,37 | 3 |
| MQQPQISVYSGSD | PHC3 | sp\|Q8NDX5\|PHC3_HUMAN | 19,22 | 6,23E+05 | 23,18 | 3 |
| NIILTTMPAGTK | NFRKB | sp\|Q6P4R8\|NFRKB_HUMAN | 19,21 | 6,06E+05 | 2,51 | 3 |
| VISGTTLGYLSPK | STAU2 | sp\|Q9NUL3\|STAU2_HUMAN | 19,08 | 5,55E+05 | 3,95 | 3 |
| SSVLSNSIPPQSST | QSER1 | sp\|Q2KHR3\|QSER1_HUMAN | 19,07 | 5,62E+05 | 24,36 | 3 |
| TQPRPTLTLSQAP | TAF6 | sp\|P49848\|TAF6_HUMAN | 19,06 | 5,54E+05 | 18,18 | 3 |
| GQPGTILRTVPM | HCFC1 | sp\|P51610\|HCFC1_HUMAN | 19,06 | 5,52E+05 | 17,50 | 3 |
| LNSPPSSIYK | FNDC3B | sp\|Q53EP0\|FND3B_HUMAN | 19,06 | 5,49E+05 | 14,62 | 3 |
| TTVVPLISTIAGDS | VEZF1 | sp\|Q14119\|VEZF1_HUMAN | 19,02 | 5,34E+05 | 9,08 | 3 |
| PSTTIENISVSVHQ | UBAP2 | \|A0A8C8KGQ8\|A0A8C8KGQ8_HUMA | 19,00 | 5,31E+05 | 17,13 | 3 |
| SFAAAPVPTTTLV | SON | sp\|P18583\|SON_HUMAN | 18,96 | 5,12E+05 | 6,27 | 3 |
| AADPPRYTFAPSV | PDLIM7 | sp\|Q9NR12\|PDLI7_HUMAN | 18,95 | 5,08E+05 | 0,11 | 3 |
| VSCTPCNIPIGTPV | NCOR1 | sp\|O75376\|NCOR1_HUMAN | 18,93 | 5,01E+05 | 7,68 | 3 |
| SPAAPIATFSLLPA | NUP214 | sp\|P35658\|NU214_HUMAN | 18,91 | 4,98E+05 | 18,48 | 3 |
| LQVTMPGIK | AHNAK | sp\|Q09666\|AHNK_HUMAN | 18,89 | 4,85E+05 | 4,88 | 3 |
| TTSSSGTNPSSSA | VEZF1 | sp\|Q14119\|VEZF1_HUMAN | 18,82 | 4,70E+05 | 21,37 | 3 |
| PAPAGFSQHPGL | SEC16A | sp\|O15027\|SC16A_HUMAN | 18,74 | 4,39E+05 | 10,72 | 3 |
| FNVPVSLESK | SARNP | sp\|P82979\|SARNP_HUMAN | 18,61 | 4,01E+05 | 2,12 | 3 |
| PPMPVLTPVHTSS | BCORL1 | sp\|Q5H9F3\|BCORL_HUMAN | 18,61 | 3,99E+05 | 1,76 | 3 |
| ATTGTPLVTMRPA | HCFC1 | sp\|P51610\|HCFC1_HUMAN | 18,59 | 3,97E+05 | 9,22 | 3 |
| FGTNPGGLFGQQ | NUP98 | \|A0A8V8TRC7\|A0A8V8TRC7_HUMA | 18,59 | 3,99E+05 | 17,29 | 3 |
| TFSTQIVR | AHNAK2 | sp\|Q8IVF2\|AHNK2_HUMAN | 18,57 | 3,88E+05 | NA | 2 |
| LTSQGAAGGSHDA | FOXK1 | sp\|P85037\|FOXK1_HUMAN | 18,56 | 4,18E+05 | 49,14 | 3 |
| ALKDTYMLSSTVS | PGM2 | sp\|Q96G03\|PGM2_HUMAN | 18,52 | 3,80E+05 | NA | 2 |
| TSSHSLQQGQQP | CNOT4 | sp\|O95628\|CNOT4_HUMAN | 18,48 | 3,68E+05 | 9,49 | 3 |
| SDTTEYQPILSSYS | RPRD2 | sp\|Q5VT52\|RPRD2_HUMAN | 18,48 | 3,66E+05 | 6,24 | 3 |
| TFGGFASSSFGEQ | NUP214 | sp\|P35658\|NU214_HUMAN | 18,47 | 3,64E+05 | 6,84 | 3 |
| QSSSLTSVPPTTFS | PRRC2C | sp\|Q9Y520\|PRC2C_HUMAN | 18,46 | 3,63E+05 | 15,47 | 3 |
| HYSPWSCGTIGSC | RC3H2 | sp\|Q9HBD1\|RC3H2_HUMAN | 18,45 | 3,71E+05 | 30,21 | 3 |
| YGARPVSSAASVY | KRT18 | sp\|P05783\|K1C18_HUMAN | 18,43 | 3,55E+05 | 13,65 | 3 |
| FTFGNSAAPAAA | POM121 | sp\|Q96HA1\|P121A_HUMAN | 18,43 | 3,52E+05 | 4,04 | 3 |
| VPASHLQQGTASG | NFRKB | sp\|Q6P4R8\|NFRKB_HUMAN | 18,37 | 3,41E+05 | 11,31 | 3 |
| PTTLPGTHPGLSET | TCF12 | sp\|Q99081\|HTF4_HUMAN | 18,37 | 3,46E+05 | 24,87 | 3 |
| QLVGQTVLNPVT | MNT | sp\|Q99583\|MNT_HUMAN | 18,34 | 3,34E+05 | 15,46 | 3 |
| QVYATMPINSFR | ZNF148 | sp\|Q9UQR1\|ZN148_HUMAN | 18,32 | 3,28E+05 | 3,56 | 3 |
| GQVSTMVTTSTTT | CNOT1 | sp\|A5YKK6\|CNOT1_HUMAN | 18,32 | 3,30E+05 | 18,61 | 3 |
| HSSPASSNYQQTT | NIPBL | sp\|Q6KC79\|NIPBL_HUMAN | 18,31 | 3,29E+05 | 17,61 | 3 |
| LATAPNVQQIQVP | LIN54 | sp\|Q6MZP7\|LIN54_HUMAN | 18,27 | 3,19E+05 | 17,51 | 3 |
| SFASSTHCQTLQN | QSER1 | sp\|Q2KHR3\|QSER1_HUMAN | 18,22 | 3,09E+05 | 19,41 | 3 |
| PTTGYSAVPANIIP | HIVEP1 | sp\|P15822\|ZEP1_HUMAN | 18,19 | 3,16E+05 | 36,23 | 3 |
| SSSLNTTLPSTSAV | TNRC6A | sp\|Q8NDV7\|TNR6A_HUMAN | 18,16 | 3,00E+05 | 28,27 | 3 |
| ILPTLPQQLQVAP | CIC | sp\|Q96RK0\|CIC_HUMAN | 18,12 | 2,86E+05 | 8,72 | 3 |
| TSPAPHLVAGPLL | CIC | sp\|Q96RK0\|CIC_HUMAN | 18,10 | 2,80E+05 | 2,93 | 3 |
| STFSFSMTKPSEK | NUP153 | sp\|P49790\|NU153_HUMAN | 18,09 | 2,81E+05 | 14,76 | 3 |
| TAPPTYTETLSTAP | SYNPO | sp\|Q8N3V7\|SYNPO_HUMAN | 18,08 | 2,79E+05 | 17,81 | 3 |
| PTPIQPTFGGATH | POM121C | sp\|A8CG34\|P121C_HUMAN | 18,06 | 2,75E+05 | 11,62 | 3 |

| | | | | | | |
|---|---|---|---|---|---|---|
| GLLPDPPRSSYLES | YLPM1 | sp\|P49750\|YLPM1_HUMAN | 18,06 | 2,76E+05 | 17,11 | 3 |
| SFYGSDIGNVVLDN | TRAK1 | sp\|Q9UPV9\|TRAK1_HUMAN | 18,04 | 2,71E+05 | 6,87 | 3 |
| VSTSAGGIMTAPAI | PICALM | sp\|Q13492\|PICAL_HUMAN | 18,01 | 2,64E+05 | 11,06 | 3 |
| YPHGVPSDSSLGHS | HIVEP2 | sp\|P31629\|ZEP2_HUMAN | 18,00 | 2,63E+05 | NA | 2 |
| FGGSPGFGGVPAF | NUP214 | sp\|P35658\|NU214_HUMAN | 18,00 | 4,26E+05 | 77,05 | 3 |
| AYALTSPLQLLATG | FOXK1 | sp\|P85037\|FOXK1_HUMAN | 17,98 | 2,61E+05 | 17,00 | 3 |
| QTTASTRPSVSAPT | SBNO1 | sp\|A3KN83\|SBNO1_HUMAN | 17,95 | 2,62E+05 | NA | 2 |
| QSPQTSFPYK | MAP7D3 | sp\|Q8IWC1\|MA7D3_HUMAN | 17,94 | 2,52E+05 | NA | 2 |
| SSNLDPSQAPSLA | UBAP2L | sp\|Q14157\|UBP2L_HUMAN | 17,87 | 2,41E+05 | 14,50 | 3 |
| TSAVPANIIPPPHPL | HIVEP1 | sp\|P15822\|ZEP1_HUMAN | 17,84 | 2,50E+05 | 46,44 | 3 |
| TTGSTYSAITTTHS | RIPOR1 | sp\|Q6ZS17\|RIPR1_HUMAN | 17,83 | 2,41E+05 | 29,42 | 3 |
| PQPLSPASSSYSVS | ATF6 | sp\|P18850\|ATF6A_HUMAN | 17,82 | 2,33E+05 | 17,23 | 3 |
| LHTSPWMPVVTTS | ZNF318 | sp\|Q5VUA4\|ZN318_HUMAN | 17,75 | 2,22E+05 | 7,78 | 3 |
| TSAPSFGFGTNTS | NUP98 | tr\|H0YEN4\|H0YEN4_HUMAN | 17,75 | 2,20E+05 | 2,34 | 3 |
| VPTTLMPVNTVALI | MAVS | sp\|Q7Z434\|MAVS_HUMAN | 17,75 | 2,22E+05 | 19,34 | 3 |
| NPPGASTYNK | HNRNPUL1 | sp\|Q9BUJ2\|HNRL1_HUMAN | 17,73 | 2,18E+05 | NA | 2 |
| NAITSSYSSTR | POM121C | sp\|A8CG34\|P121C_HUMAN | 17,72 | 2,18E+05 | 18,75 | 3 |
| ISSLNPAYSQYSQK | HIVEP2 | sp\|P31629\|ZEP2_HUMAN | 17,71 | 2,15E+05 | 10,59 | 3 |
| LPNPTKPNNVPSV | ATF7IP | sp\|Q6VMQ6\|MCAF1_HUMAN | 17,69 | 2,13E+05 | 13,08 | 3 |
| SKPEPPAMPQPVP | RPS3 | sp\|P23396\|RS3_HUMAN | 17,66 | 2,07E+05 | 9,93 | 3 |
| LSTMSSSLYASSQL | PPP1R12A | sp\|O14974\|MYPT1_HUMAN | 17,65 | 2,07E+05 | NA | 2 |
| VPVTLPERK | HIVEP3 | sp\|Q5T1R4\|ZEP3_HUMAN | 17,61 | 2,01E+05 | 7,08 | 3 |
| LMSMPVMTVDYSI | SETD2 | sp\|Q9BYW2\|SETD2_HUMAN | 17,61 | 2,06E+05 | 27,38 | 3 |
| NPIMVTLAPNIQTG | TAB2 | sp\|Q9NYJ8\|TAB2_HUMAN | 17,61 | 2,01E+05 | 11,78 | 3 |
| QTLPTSNYFTTVSE | KDM3B | sp\|Q7LBC6\|KDM3B_HUMAN | 17,61 | 2,07E+05 | 30,07 | 3 |
| QALGSESFLPGSSFA | MAMLD1 | sp\|Q13495\|MAMD1_HUMAN | 17,61 | 2,01E+05 | 12,82 | 3 |
| YLPNSDPLHQSDT | ANKRD17 | sp\|O75179\|ANR17_HUMAN | 17,59 | 2,01E+05 | 25,89 | 3 |
| EAIIPGSVYDR | UPF1 | sp\|Q92900\|RENT1_HUMAN | 17,58 | 1,95E+05 | 3,53 | 3 |
| TTTHSWVSG | FOXO1 | sp\|Q12778\|FOXO1_HUMAN | 17,57 | 1,95E+05 | 13,50 | 3 |
| VVPSSFNFGGPAP | IPO7 | sp\|O95373\|IPO7_HUMAN | 17,56 | 1,93E+05 | 3,30 | 3 |
| PQSIIPPVQTLSYSK | QSER1 | sp\|Q2KHR3\|QSER1_HUMAN | 17,55 | 1,94E+05 | 19,72 | 3 |
| VPMSTAGQTSR | SH3RF1 | sp\|Q7Z6J0\|SH3R1_HUMAN | 17,54 | 1,92E+05 | 11,85 | 3 |
| QLADVPGGPLFNG | ANKHD1 | sp\|Q8IWZ3\|ANKH1_HUMAN | 17,49 | 1,90E+05 | 26,49 | 3 |
| GHGFQFVSLSSPLH | ZNF281 | sp\|Q9Y2X9\|ZN281_HUMAN | 17,49 | 1,84E+05 | NA | 2 |
| TSPGSVTNTSLAHE | TNRC6A | sp\|Q8NDV7\|TNR6A_HUMAN | 17,46 | 1,84E+05 | 23,24 | 3 |
| AAVAAAAQPQPSY | ZFR | sp\|Q96KR1\|ZFR_HUMAN | 17,46 | 1,81E+05 | 13,01 | 3 |
| TSQQPAGHAIHVV | FOXK1 | sp\|P85037\|FOXK1_HUMAN | 17,43 | 1,78E+05 | 12,76 | 3 |
| QTPVNTVSSTNLV | ATF7IP | sp\|Q6VMQ6\|MCAF1_HUMAN | 17,43 | 1,80E+05 | 22,73 | 3 |
| VPVTLPER | HIVEP3 | sp\|Q5T1R4\|ZEP3_HUMAN | 17,40 | 1,74E+05 | 11,58 | 3 |
| AMQVPVVMTSLG | ELF2 | sp\|Q15723\|ELF2_HUMAN | 17,39 | 1,72E+05 | NA | 2 |
| PTTTSGGLIMTSDT | CDK8 | sp\|P49336\|CDK8_HUMAN | 17,38 | 1,73E+05 | 18,84 | 3 |
| TSITTAPAATTAATS | PHACTR4 | sp\|Q8IZ21\|PHAR4_HUMAN | 17,38 | 1,73E+05 | 24,13 | 3 |
| SAYNSYSWGAN | UBAP2L | sp\|Q14157\|UBP2L_HUMAN | 17,36 | 1,70E+05 | 15,89 | 3 |
| FQVTTTANK | WNK1 | sp\|Q9H4A3\|WNK1_HUMAN | 17,36 | 1,69E+05 | NA | 2 |
| GAGFGTALGAGQ | NUP98 | r\|A0A8V8TRC7\|A0A8V8TRC7_HUMA | 17,34 | 1,66E+05 | 4,19 | 3 |
| DPVTPPSDPSIPIPT | SPEN | sp\|Q96T58\|MINT_HUMAN | 17,30 | 1,66E+05 | 25,55 | 3 |
| APVTLTSGILMGAK | TRAK1 | sp\|Q9UPV9\|TRAK1_HUMAN | 17,30 | 1,62E+05 | 6,84 | 3 |
| CVCEPGWSGPR | NOTCH3 | sp\|Q9UM47\|NOTC3_HUMAN | 17,29 | 1,61E+05 | 11,46 | 3 |
| IATTVVTTVYQEPII | OGA | sp\|O60502\|OGA_HUMAN | 17,27 | 1,59E+05 | 13,73 | 3 |
| SFGTTSGGLFGFG | NUP98 | r\|A0A8V8TRC7\|A0A8V8TRC7_HUMA | 17,27 | 1,63E+05 | 29,28 | 3 |

| | | | | | | |
|---|---|---|---|---|---|---|
| LHVQSQTQPVSLA | ZYX | sp\|Q15942\|ZYX_HUMAN | 17,24 | 1,55E+05 | 11,01 | 3 |
| HTNWTGHGGTVS: | PGD | sp\|P52209\|6PGD_HUMAN | 17,22 | 1,53E+05 | NA | 2 |
| NASSIPTVHTSPQL | KIAA2026 | sp\|Q5HYC2\|K2026_HUMAN | 17,20 | 1,51E+05 | 9,47 | 3 |
| GLPTTLGPAMVTF | MTSS1 | sp\|O43312\|MTSS1_HUMAN | 17,18 | 1,49E+05 | NA | 2 |
| NVLPPSSIGFTFSV | NUP153 | sp\|P49790\|NU153_HUMAN | 17,18 | 1,56E+05 | 39,23 | 3 |
| TVTTTAGNVISVIG: | RC3H2 | sp\|Q9HBD1\|RC3H2_HUMAN | 17,16 | 1,53E+05 | 33,11 | 3 |
| PQTVPSGMAGPPF | SEC16A | sp\|O15027\|SC16A_HUMAN | 17,15 | 1,48E+05 | 22,83 | 3 |
| ITLPSHPALGTPK | SAP130 | sp\|Q9H0E3\|SP130_HUMAN | 17,14 | 1,47E+05 | 24,22 | 3 |
| GSVTTLK | SPEN | sp\|Q96T58\|MINT_HUMAN | 17,12 | 1,43E+05 | NA | 2 |
| RLPDAHSDYAR | RBM14 | sp\|Q96PK6\|RBM14_HUMAN | 17,09 | 1,40E+05 | 4,27 | 3 |
| SQMAPGVPGHTV | FOXK1 | sp\|P85037\|FOXK1_HUMAN | 17,07 | 1,55E+05 | NA | 2 |
| AVPVTLASQQAGT | ARID3B | sp\|Q8IVW6\|ARI3B_HUMAN | 17,06 | 1,37E+05 | 11,01 | 3 |
| ATTTSYSPSVPVSFI | SIX4 | sp\|Q9UIU6\|SIX4_HUMAN | 17,05 | 1,37E+05 | 14,26 | 3 |
| CLTGFTGQK | NOTCH2 | sp\|Q04721\|NOTC2_HUMAN | 17,02 | 1,33E+05 | NA | 2 |
| LASTVSQSNVVIAF | MLXIP | sp\|Q9HAP2\|MLXIP_HUMAN | 17,00 | 1,31E+05 | 3,98 | 3 |
| LGATPGGKPTAIHC | KANSL3 | sp\|Q9P2N6\|KANL3_HUMAN | 16,99 | 1,31E+05 | 11,94 | 3 |
| /SVYQEPEGSGLDI | FGD5 | sp\|Q6ZNL6\|FGD5_HUMAN | 16,98 | 1,33E+05 | NA | 2 |
| PFTTGSQDVSNAF | SEC23IP | sp\|Q9Y6Y8\|S23IP_HUMAN | 16,96 | 1,30E+05 | 23,04 | 3 |
| AQPETPVTLQFQG | EP400 | sp\|Q96L91\|EP400_HUMAN | 16,94 | 1,27E+05 | 16,87 | 3 |
| MTVLSTAQSDYNR | HIVEP1 | sp\|P15822\|ZEP1_HUMAN | 16,94 | 1,26E+05 | 7,30 | 3 |
| PSTLHFPTSPIIQQP | NFIA | tr\|S4R3W2\|S4R3W2_HUMAN | 16,90 | 1,24E+05 | NA | 2 |
| SPVLSPTLPAEAPT | DCP1A | sp\|Q9NPI6\|DCP1A_HUMAN | 16,89 | 1,22E+05 | NA | 2 |
| LVQLLTTTAEQQLF | NCOA1 | sp\|Q15788\|NCOA1_HUMAN | 16,88 | 1,21E+05 | 7,10 | 3 |
| AVPTSIYQTSTGQY | CREM | tr\|C9IYM9\|C9IYM9_HUMAN | 16,88 | 1,21E+05 | NA | 2 |
| MTPSDSCEDTQMT | HIVEP2 | sp\|P31629\|ZEP2_HUMAN | 16,86 | 1,19E+05 | 9,91 | 3 |
| CSFHPIPTR | JMJD1C | sp\|Q15652\|JHD2C_HUMAN | 16,85 | 1,19E+05 | 15,44 | 3 |
| /DAPASAFGSSLLG | PPP1R13L | sp\|Q8WUF5\|IASPP_HUMAN | 16,85 | 1,19E+05 | 13,49 | 3 |
| QSAAVTPSSTTSST | ADRM1 | sp\|Q16186\|ADRM1_HUMAN | 16,85 | 1,19E+05 | 16,34 | 3 |
| VSTPATTTSTFSR | SYNPO | sp\|Q8N3V7\|SYNPO_HUMAN | 16,80 | 1,16E+05 | 23,16 | 3 |
| VTSSSFFAAPNPY\ | ERG | sp\|P11308\|ERG_HUMAN | 16,80 | 1,17E+05 | 26,58 | 3 |
| ATPSTYSGVFR | PRRC2A | sp\|P48634\|PRC2A_HUMAN | 16,79 | 1,15E+05 | NA | 2 |
| TPSPVSVSMKPDLF | SPEN | sp\|Q96T58\|MINT_HUMAN | 16,78 | 1,13E+05 | NA | 2 |
| /YNPSHSDSLASQ( | SEC16A | sp\|O15027\|SC16A_HUMAN | 16,78 | 1,13E+05 | 4,78 | 3 |
| HSHAVSTAAMTR | HCFC1 | sp\|P51610\|HCFC1_HUMAN | 16,77 | 1,44E+05 | 84,40 | 3 |
| GTGVINSTPAPAN | NUP153 | sp\|P49790\|NU153_HUMAN | 16,77 | 1,13E+05 | NA | 2 |
| HLCPVTLDGER | ZNF407 | sp\|Q9C0G0\|ZN407_HUMAN | 16,77 | 1,12E+05 | 10,80 | 3 |
| VPSQSSVVYHPYG | HOXC9 | sp\|P31274\|HXC9_HUMAN | 16,77 | 1,34E+05 | 59,11 | 3 |
| QTYASMVTSSHLF | NFATC3 | sp\|Q12968\|NFAC3_HUMAN | 16,76 | 1,12E+05 | 13,50 | 3 |
| VGFNEMEAPTTAY | HCLS1 | sp\|P14317\|HCLS1_HUMAN | 16,75 | 1,11E+05 | 9,13 | 3 |
| GGAGGSPSVTWA | BICRA | sp\|Q9NZM4\|BICRA_HUMAN | 16,74 | 1,10E+05 | 10,86 | 3 |
| LPVIMGSFAAPVCT | TRAK2 | sp\|O60296\|TRAK2_HUMAN | 16,67 | 1,06E+05 | 17,68 | 3 |
| HQVVYTTLPAPPA( | ATF7IP | sp\|Q6VMQ6\|MCAF1_HUMAN | 16,67 | 1,05E+05 | 16,75 | 3 |
| TGLPTVPSSAYSHF | ALMS1 | sp\|Q8TCU4\|ALMS1_HUMAN | 16,65 | 1,03E+05 | 12,21 | 3 |
| SGNPSHGTLGLSG | PROSER1 | sp\|Q86XN7\|PRSR1_HUMAN | 16,63 | 1,03E+05 | 24,40 | 3 |
| IESSQYIGPDMLPN | HIVEP2 | sp\|P31629\|ZEP2_HUMAN | 16,62 | 1,01E+05 | NA | 2 |
| QGPVGVNVTYGG | FLNA | sp\|P21333\|FLNA_HUMAN | 16,61 | 1,01E+05 | 10,92 | 3 |
| PLTSFGSAPSSEGAI | PRRC2C | sp\|Q9Y520\|PRC2C_HUMAN | 16,60 | 9,95E+04 | 9,67 | 3 |
| AHSPASLSFASYR | PARP4 | sp\|Q9UKK3\|PARP4_HUMAN | 16,60 | 1,11E+05 | 49,84 | 3 |
| SLYASSPGGVYATF | VIM | sp\|P08670\|VIME_HUMAN | 16,60 | 1,01E+05 | 23,96 | 3 |
| TPPMPVLTPVHTS: | BCORL1 | sp\|Q5H9F3\|BCORL_HUMAN | 16,58 | 9,82E+04 | NA | 2 |

| | | | | | | |
|---|---|---|---|---|---|---|
| LHTASSTMYSNTNI | HDX | sp\|Q7Z353\|HDX_HUMAN | 16,56 | 9,71E+04 | 7,81 | 3 |
| ATMEGIGAIGGTPI | NONO | sp\|Q15233\|NONO_HUMAN | 16,55 | 9,68E+04 | 14,94 | 3 |
| PKPQSIIPPVQTLSY | QSER1 | sp\|Q2KHR3\|QSER1_HUMAN | 16,54 | 9,80E+04 | 26,65 | 3 |
| TVDSISVNTSLDQN | TNRC6A | sp\|Q8NDV7\|TNR6A_HUMAN | 16,54 | 9,75E+04 | 26,81 | 3 |
| GNPTFAAVTAGYD | TMEM131 | sp\|Q92545\|TM131_HUMAN | 16,54 | 9,56E+04 | 9,70 | 3 |
| GNVEPASLPSASV | NUP153 | sp\|P49790\|NU153_HUMAN | 16,53 | 9,55E+04 | 18,44 | 3 |
| ISSSSFIEPIFPTSK | TMTC3 | sp\|Q6ZXV5\|TMTC3_HUMAN | 16,50 | 9,30E+04 | 7,33 | 3 |
| VRSIYDDISSPGLGS | NUP35 | sp\|Q8NFH5\|NUP35_HUMAN | 16,49 | 9,24E+04 | NA | 2 |
| TLAQHISEVITQDY | NCOR2 | sp\|Q9Y618\|NCOR2_HUMAN | 16,48 | 9,17E+04 | 6,91 | 3 |
| SGYVPPPVATPFS | ZYX | sp\|Q15942\|ZYX_HUMAN | 16,46 | 9,05E+04 | NA | 2 |
| LMLSTSEYSQSPK | TP53BP1 | sp\|Q12888\|TP53B_HUMAN | 16,45 | 9,01E+04 | 14,90 | 3 |
| VLVETSYPSQTTR | EGR1 | sp\|P18146\|EGR1_HUMAN | 16,44 | 9,44E+04 | NA | 2 |
| GGGLLKPTVASQN | PICALM | tr\|H0YEF7\|H0YEF7_HUMAN | 16,44 | 8,88E+04 | NA | 2 |
| FTPSSTMMEVFLC | UBAP2L | sp\|Q14157\|UBP2L_HUMAN | 16,43 | 8,87E+04 | 12,86 | 3 |
| QQTSAVNGRPLPI | FOXO1 | sp\|Q12778\|FOXO1_HUMAN | 16,41 | 8,70E+04 | NA | 2 |
| KKPLFITTDSSK | KDM3B | sp\|Q7LBC6\|KDM3B_HUMAN | 16,40 | 8,70E+04 | 12,62 | 3 |
| LEGPSATAPASSPG | CC2D1A | sp\|Q6P1N0\|C2D1A_HUMAN | 16,40 | 8,72E+04 | 17,03 | 3 |
| APGPGPSTFSFVPP | NUP214 | sp\|P35658\|NU214_HUMAN | 16,39 | 8,60E+04 | NA | 2 |
| SFPSPPAVSIASFV | POGZ | sp\|Q7Z3K3\|POGZ_HUMAN | 16,36 | 1,00E+05 | 65,31 | 3 |
| ASQAGMTGYGMI | TAGLN2 | sp\|P37802\|TAGL2_HUMAN | 16,33 | 8,27E+04 | 9,09 | 3 |
| EVVKPVPITSPAVSI | PDLIM5 | sp\|Q96HC4\|PDLI5_HUMAN | 16,32 | 9,14E+04 | NA | 2 |
| PVTPPSDPSIPIPTL | SPEN | sp\|Q96T58\|MINT_HUMAN | 16,32 | 8,16E+04 | NA | 2 |
| IVSFGAVGFDPHPI | TLE3 | sp\|Q04726\|TLE3_HUMAN | 16,30 | 8,10E+04 | 9,76 | 3 |
| WPQPVTTSHYAHD | TNS1 | sp\|Q9HBL0\|TENS1_HUMAN | 16,28 | 8,16E+04 | NA | 2 |
| IDDTPRGSWACSII | DOCK7 | sp\|Q96N67\|DOCK7_HUMAN | 16,28 | 7,95E+04 | NA | 2 |
| FNFGIITPTPSSNFT | NUP214 | sp\|P35658\|NU214_HUMAN | 16,27 | 7,99E+04 | NA | 2 |
| AVGFNEMEAPTTA | HCLS1 | sp\|P14317\|HCLS1_HUMAN | 16,27 | 8,04E+04 | 22,62 | 3 |
| NLRVTEANGR | ZHX3 | sp\|Q9H4I2\|ZHX3_HUMAN | 16,27 | 7,90E+04 | NA | 2 |
| DGSTQVTVEEPVC | PLIN3 | sp\|O60664\|PLIN3_HUMAN | 16,26 | 7,97E+04 | 18,63 | 3 |
| LPQSNYFTTLSNSV | JMJD1C | sp\|Q15652\|JHD2C_HUMAN | 16,26 | 8,20E+04 | 36,42 | 3 |
| MPQQIGIPTSSLTQ | WNK1 | sp\|Q9H4A3\|WNK1_HUMAN | 16,25 | 7,94E+04 | 20,82 | 3 |
| NQPASFAVSLNGA | FLNA | sp\|P21333\|FLNA_HUMAN | 16,24 | 7,76E+04 | 2,80 | 3 |
| VPQQLQVHGVQC | RFX1 | sp\|P22670\|RFX1_HUMAN | 16,23 | 7,75E+04 | 18,46 | 3 |
| NVATHSTLVLTAQT | SPEN | sp\|Q96T58\|MINT_HUMAN | 16,22 | 7,63E+04 | NA | 2 |
| IMQSIANSLPPHM | EMSY | sp\|Q7Z589\|EMSY_HUMAN | 16,20 | 8,37E+04 | NA | 2 |
| YVMTTTTLER | EIF2S1 | sp\|P05198\|IF2A_HUMAN | 16,20 | 7,62E+04 | NA | 2 |
| REDENAEPVGTTYC | DBN1 | sp\|Q16643\|DREB_HUMAN | 16,17 | 7,37E+04 | 5,45 | 3 |
| NYNVPLSSTAQSTS | TNRC6A | sp\|Q8NDV7\|TNR6A_HUMAN | 16,16 | 7,42E+04 | 18,02 | 3 |
| SGYWASHLAGRK | UGGT1 | sp\|Q9NYU2\|UGGG1_HUMAN | 16,16 | 7,32E+04 | 5,61 | 3 |
| TETVTPSEAPVLAA | TXNDC5 | sp\|Q8NBS9\|TXND5_HUMAN | 16,13 | 7,39E+04 | 30,71 | 3 |
| YGDSLYYNNAYGA | RBM4 | sp\|Q9BWF3\|RBM4_HUMAN | 16,10 | 8,90E+04 | 65,00 | 3 |
| WEVLSATPTTIK | SP3 | sp\|Q02447\|SP3_HUMAN | 16,09 | 7,40E+04 | NA | 2 |
| LGHHPPVTIGQPQ | HELZ | sp\|P42694\|HELZ_HUMAN | 16,08 | 6,95E+04 | NA | 2 |
| ATAAAYGGYPTAH | ZFR | sp\|Q96KR1\|ZFR_HUMAN | 16,08 | 7,42E+04 | 46,92 | 3 |
| GISAPQVSIPDVNY | AHNAK | sp\|Q09666\|AHNK_HUMAN | 16,08 | 6,94E+04 | 9,07 | 3 |
| YDDISSPGLGSTPLT | NUP35 | sp\|Q8NFH5\|NUP35_HUMAN | 16,07 | 6,93E+04 | 13,58 | 3 |
| AQPTILTTTATLPA | SAP30BP | sp\|Q9UHR5\|S30BP_HUMAN | 16,06 | 6,88E+04 | NA | 2 |
| FDELGGLLKPTVAS | PICALM | sp\|Q13492\|PICAL_HUMAN | 16,06 | 6,82E+04 | 2,41 | 3 |
| PSSTWGASPLGWT | TNRC6C | sp\|Q9HCJ0\|TNR6C_HUMAN | 16,05 | 6,89E+04 | 21,90 | 3 |
| TPGCNPQLTYTATL | USP54 | sp\|Q70EL1\|UBP54_HUMAN | 16,04 | 6,79E+04 | 12,16 | 3 |

| | | | | | | |
|---|---|---|---|---|---|---|
| DYTTPSSSLPNTVAT | PRRC2C | sp\|Q9Y520\|PRC2C_HUMAN | 16,04 | 6,88E+04 | 25,32 | 3 |
| LDVGPQAPVGSVY | DBNL | sp\|Q9UJU6\|DBNL_HUMAN | 16,03 | 6,73E+04 | 12,06 | 3 |
| NMQPLYVLQTLPI | KMT2A | sp\|Q03164\|KMT2A_HUMAN | 16,01 | 6,61E+04 | 4,94 | 3 |
| SGPTSHTQASLSH | TNRC6C | sp\|Q9HCJ0\|TNR6C_HUMAN | 16,00 | 6,57E+04 | NA | 2 |
| VMTSGTGAPAK | HCFC1 | sp\|P51610\|HCFC1_HUMAN | 16,00 | 6,58E+04 | NA | 2 |
| GSWACSIFDLK | DOCK7 | sp\|Q96N67\|DOCK7_HUMAN | 15,99 | 6,59E+04 | NA | 2 |
| GIDYQAGDTYVST | CDK13 | sp\|Q14004\|CDK13_HUMAN | 15,98 | 6,47E+04 | 8,68 | 3 |
| NASASTISPPSSME | MAP1B | sp\|P46821\|MAP1B_HUMAN | 15,97 | 6,65E+04 | 34,19 | 3 |
| VFLLSVPSLDCLPI | HIVEP1 | sp\|P15822\|ZEP1_HUMAN | 15,96 | 6,56E+04 | 26,48 | 3 |
| TDEAQSNLNTTVG | ETAA1 | sp\|Q9NY74\|ETAA1_HUMAN | 15,94 | 6,48E+04 | 29,30 | 3 |
| SFSSMSLPGAPTA | ARNT | sp\|P27540\|ARNT_HUMAN | 15,92 | 6,27E+04 | 17,27 | 3 |
| IGSSPGVMGTSVA | NUP214 | sp\|P35658\|NU214_HUMAN | 15,90 | 6,14E+04 | 7,99 | 3 |
| AVTTVPSMGIGLV | TMEM263 | sp\|Q8WUH6\|TM263_HUMAN | 15,89 | 6,08E+04 | 3,59 | 3 |
| IGSFAEPSSVSFSSK | KMT2A | sp\|Q03164\|KMT2A_HUMAN | 15,89 | 6,06E+04 | NA | 2 |
| PSSFSSMSLPGAPT | ARNT | sp\|P27540\|ARNT_HUMAN | 15,88 | 6,06E+04 | NA | 2 |
| FPQHSQSFPTSSTY | RAI1 | sp\|Q7Z5J4\|RAI1_HUMAN | 15,87 | 6,00E+04 | NA | 2 |
| TIPGTYVGVANPVP | BCORL1 | sp\|Q5H9F3\|BCORL_HUMAN | 15,87 | 6,00E+04 | 4,69 | 3 |
| SFGQFTPQSSQGT | SETD1A | sp\|O15047\|SET1A_HUMAN | 15,87 | 7,71E+04 | 67,03 | 3 |
| AQTPVTISANQIIL | LIN54 | sp\|Q6MZP7\|LIN54_HUMAN | 15,80 | 5,82E+04 | 23,97 | 3 |
| LVAFSAVAEAVSSY | TET3 | sp\|O43151\|TET3_HUMAN | 15,80 | 5,76E+04 | NA | 2 |
| LLQTPIVTNHLVQLY | HROB | sp\|Q8N3J3\|HROB_HUMAN | 15,78 | 5,64E+04 | NA | 2 |
| AFGAQTSTTADQG | NUP153 | sp\|P49790\|NU153_HUMAN | 15,75 | 5,82E+04 | 41,80 | 3 |
| TQFTTPLASFTTVA | BRD8 | sp\|Q9H0E9\|BRD8_HUMAN | 15,74 | 5,51E+04 | NA | 2 |
| YLPSQPSSLPPSYFG | SEC23IP | sp\|Q9Y6Y8\|S23IP_HUMAN | 15,73 | 5,73E+04 | NA | 2 |
| APSAVVYTVPNTGC | SIX4 | sp\|Q9UIU6\|SIX4_HUMAN | 15,66 | 5,21E+04 | NA | 2 |
| FTFGNSAAPAPATA | POM121C | sp\|A8CG34\|P121C_HUMAN | 15,65 | 5,55E+04 | 44,25 | 3 |
| LVTATTTAVSSVLL | DCP1A | sp\|Q9NPI6\|DCP1A_HUMAN | 15,64 | 5,24E+04 | 26,76 | 3 |
| TANVPQTVPMR | YAP1 | sp\|P46937\|YAP1_HUMAN | 15,64 | 5,13E+04 | NA | 2 |
| QQSYLDSGIHSGAT | CTNNB1 | sp\|P35222\|CTNB1_HUMAN | 15,64 | 5,14E+04 | NA | 2 |
| PLTPPVVVTHGVC | SPEN | sp\|Q96T58\|MINT_HUMAN | 15,63 | 5,24E+04 | 27,33 | 3 |
| YASYSGTVTQATFT | TASOR2 | sp\|Q5VWN6\|TASO2_HUMAN | 15,61 | 5,03E+04 | NA | 2 |
| STFSFSMTK | NUP153 | sp\|P49790\|NU153_HUMAN | 15,60 | 4,99E+04 | NA | 2 |
| TMSQGGMVTVIP | TOX4 | sp\|O94842\|TOX4_HUMAN | 15,60 | 4,96E+04 | 5,63 | 3 |
| TSPSGLDPNASVYC | MACO1 | sp\|Q8N5G2\|MACOI_HUMAN | 15,58 | 4,90E+04 | NA | 2 |
| SQPQSSMGYSATS | CDK8 | sp\|P49336\|CDK8_HUMAN | 15,55 | 4,91E+04 | NA | 2 |
| TMAVPTSIYQTSTG | CREM | tr\|C9IYM9\|C9IYM9_HUMAN | 15,49 | 5,82E+04 | NA | 2 |
| AAEPHVVTVTMN | BMPR2 | sp\|Q13873\|BMPR2_HUMAN | 15,47 | 4,56E+04 | NA | 2 |
| YGVVSTYDSSLSSY | ATXN2 | sp\|Q99700\|ATX2_HUMAN | 15,46 | 4,52E+04 | NA | 2 |
| TVYCQASFPGANIIC | NR3C1 | sp\|P04150\|GCR_HUMAN | 15,46 | 4,54E+04 | 17,64 | 3 |
| LDLTGTAATATSFA | FOXP4 | sp\|Q8IVH2\|FOXP4_HUMAN | 15,45 | 4,47E+04 | 4,49 | 3 |
| SSSTTGLPDMTGS | UBAP2 | \|A0A8C8KGQ8\|A0A8C8KGQ8_HUMA | 15,44 | 4,46E+04 | NA | 2 |
| TLHVTSNPVHAAD | NFRKB | sp\|Q6P4R8\|NFRKB_HUMAN | 15,37 | 4,49E+04 | NA | 2 |
| SSAASVYAGAGGSC | KRT18 | sp\|P05783\|K1C18_HUMAN | 15,36 | 4,27E+04 | 19,60 | 3 |
| GGSGGGSFGDNLV | LMNA | sp\|P02545\|LMNA_HUMAN | 15,36 | 4,25E+04 | 15,04 | 3 |
| IGQVTMPMVMPS | RNF214 | sp\|Q8ND24\|RN214_HUMAN | 15,34 | 4,16E+04 | NA | 2 |
| RPGPGQTGLTVTS | NFRKB | sp\|Q6P4R8\|NFRKB_HUMAN | 15,31 | 4,06E+04 | NA | 2 |
| GPVLATSSGAGVF | WNK1 | sp\|Q9H4A3\|WNK1_HUMAN | 15,29 | 4,03E+04 | 14,19 | 3 |
| YGGMSQYNCAPGI | FOXO1 | sp\|Q12778\|FOXO1_HUMAN | 15,28 | 4,00E+04 | NA | 2 |
| STEIGTMISVSSAEY | PRRC2C | sp\|Q9Y520\|PRC2C_HUMAN | 15,26 | 4,18E+04 | 45,57 | 3 |
| AVTLAVTSVAGSPA | MTCL1 | sp\|Q9Y4B5\|MTCL1_HUMAN | 15,23 | 4,02E+04 | 32,73 | 3 |

| | | | | | | |
|---|---|---|---|---|---|---|
| VNRTTLTITQPRPT | TAF6 | sp\|P49848\|TAF6_HUMAN | 15,22 | 3,85E+04 | NA | 2 |
| GNFLHSFQNDNYSF | SSBP3 | sp\|Q9BWW4\|SSBP3_HUMAN | 15,20 | 5,01E+04 | 74,60 | 3 |
| NPTQGATPTWTPT | ARNT | sp\|P27540\|ARNT_HUMAN | 15,14 | 4,39E+04 | NA | 2 |
| SSCTTVYNGYGK | FNDC3B | sp\|Q53EP0\|FND3B_HUMAN | 15,04 | 3,39E+04 | NA | 2 |
| VLSYEGGMSVTQ( | NCOR2 | sp\|Q9Y618\|NCOR2_HUMAN | 15,02 | 3,91E+04 | 56,50 | 3 |
| GIDYQAGDTYVSTS | CDK13 | sp\|Q14004\|CDK13_HUMAN | 14,93 | 3,15E+04 | 14,08 | 3 |
| SSLQDPVSTDEDVI | HIVEP1 | sp\|P15822\|ZEP1_HUMAN | 14,89 | 3,04E+04 | NA | 2 |
| QQTLDTTSSVPAP | UBAP2 | \|A0A8C8KGQ8\|A0A8C8KGQ8_HUMA | 14,86 | 3,07E+04 | 29,67 | 3 |
| ESGVIGSTLNSTTQ | LIN54 | sp\|Q6MZP7\|LIN54_HUMAN | 14,76 | 2,78E+04 | NA | 2 |
| PTFADAYSNMGNT | OGT | sp\|O15294\|OGT1_HUMAN | 14,73 | 2,94E+04 | 46,90 | 3 |
| ALYTVPTQNVTPN | SEC24B | sp\|O95487\|SC24B_HUMAN | 14,51 | 3,06E+04 | NA | 2 |
| PTPAPSTSYSPQAD | TJP1 | sp\|Q07157\|ZO1_HUMAN | 14,47 | 2,29E+04 | NA | 2 |
| SCSTEQPLTSTK | QSER1 | sp\|Q2KHR3\|QSER1_HUMAN | 13,72 | 1,41E+04 | NA | 2 |

ts
