## Supplementary material for "Mapping the endothelial O-GlcNAcome uncovers CCAR1 as a regulator of senescence": Suppl Table 2

| Gene | Protein | Thia_non | Untr_nonz | IgG_nonz | Thia_mean | Thia_CV | Untr_mean | Untr_CV |
|---|---|---|---|---|---|---|---|---|
| NUP153 | sp\|P49790\|NU | 3 | 3 | 0 | 2,69E+07 | 19,67 | 2,42E+07 | 71,10 |
| UBAP2 | sp\|Q5T6F2\|UI | 3 | 3 | 0 | 1,26E+07 | 6,87 | 1,07E+07 | 41,52 |
| PRRC2C | tr\|E7EPN9\|E7 | 3 | 3 | 0 | 8,86E+06 | 10,71 | 7,29E+06 | 53,31 |
| UBAP2L | sp\|Q14157\|UE | 3 | 3 | 0 | 8,79E+06 | 47,31 | 7,64E+06 | 61,82 |
| NUP54 | sp\|Q7Z3B4\|NI | 3 | 3 | 0 | 8,54E+06 | 14,37 | 6,90E+06 | 38,36 |
| WNK1 | sp\|Q9H4A3\|W | 3 | 3 | 0 | 8,50E+06 | 44,58 | 7,83E+06 | 32,96 |
| NUP98 | sp\|P52948\|NU | 3 | 3 | 0 | 8,10E+06 | 33,37 | 4,91E+06 | 53,42 |
| SEC24B | sp\|O95487\|SC | 3 | 3 | 0 | 7,11E+06 | 25,97 | 2,29E+06 | 43,12 |
| SEC23IP | sp\|Q9Y6Y8\|S2 | 3 | 3 | 0 | 6,52E+06 | 36,31 | 3,05E+06 | 89,09 |
| NUP88 | sp\|Q99567\|NU | 3 | 3 | 0 | 6,50E+06 | 34,48 | 4,77E+06 | 46,45 |
| SEC23A | sp\|Q15436\|SC | 3 | 3 | 0 | 5,56E+06 | 49,88 | 1,54E+06 | 47,78 |
| CCAR1 | sp\|Q8IX12\|CC | 3 | 3 | 0 | 4,58E+06 | 10,35 | 4,20E+05 | 104,58 |
| NUP62 | sp\|P37198\|NU | 3 | 3 | 0 | 4,31E+06 | 45,48 | 4,12E+06 | 53,30 |
| RAE1 | sp\|P78406\|RA | 3 | 3 | 0 | 4,09E+06 | 49,76 | 2,92E+06 | 39,65 |
| NUP58 | sp\|Q9BVL2\|N | 3 | 3 | 0 | 3,68E+06 | 15,87 | 3,24E+06 | 37,12 |
| SEC24C | sp\|P53992\|SC | 3 | 3 | 0 | 3,01E+06 | 44,33 | 1,13E+05 | 87,15 |
| CIC | tr\|A0A7P0T9K | 3 | 3 | 0 | 2,42E+06 | 30,08 | 1,38E+06 | 90,32 |
| POM121C | sp\|A8CG34\|P | 3 | 3 | 0 | 2,15E+06 | 38,31 | 1,93E+06 | 117,41 |
| ANKRD17 | sp\|O75179\|AN | 3 | 3 | 0 | 2,07E+06 | 27,96 | 8,25E+05 | 86,58 |
| RUVBL2 | sp\|Q9Y230\|RU | 3 | 3 | 0 | 1,53E+06 | 56,03 | 6,87E+05 | 43,62 |
| MEF2D | sp\|Q14814\|MI | 3 | 3 | 0 | 1,48E+06 | 49,99 | 1,03E+06 | 60,94 |
| QSER1 | tr\|A0A3B3IRW | 3 | 3 | 0 | 1,44E+06 | 46,41 | 5,34E+05 | 76,36 |
| EIF4G1 | sp\|Q04637\|IF4 | 3 | 1 | 0 | 1,43E+06 | 111,06 | 1,77E+04 | NA |
| HCFC1 | sp\|P51610\|HC | 3 | 3 | 0 | 1,36E+06 | 12,52 | 8,56E+05 | 65,67 |
| EMSY | sp\|Q7Z589\|EN | 3 | 3 | 0 | 1,26E+06 | 56,63 | 8,63E+05 | 82,20 |
| SS18 | sp\|Q15532\|SS | 3 | 3 | 0 | 1,19E+06 | 56,66 | 6,34E+05 | 68,82 |
| JMJD1C | sp\|Q15652\|JH | 3 | 3 | 0 | 1,18E+06 | 28,98 | 1,13E+06 | 82,94 |
| SEC31A | tr\|D6REX3\|D6 | 3 | 3 | 0 | 1,12E+06 | 37,37 | 5,11E+05 | 56,23 |
| SPEN | sp\|Q96T58\|MI | 3 | 3 | 0 | 1,11E+06 | 57,16 | 5,80E+05 | 150,55 |
| SETD1A | sp\|O15047\|SE | 3 | 3 | 0 | 9,47E+05 | 12,69 | 6,40E+05 | 45,60 |
| ZFR | sp\|Q96KR1\|ZI | 3 | 3 | 0 | 9,12E+05 | 54,42 | 1,88E+05 | 115,57 |
| PRRC1 | sp\|Q96M27\|P | 3 | 3 | 0 | 9,09E+05 | 21,17 | 1,13E+05 | 101,11 |
| RUVBL1 | sp\|Q9Y265\|RU | 3 | 3 | 0 | 9,07E+05 | 41,97 | 4,43E+05 | 44,42 |
| ERG | sp\|P11308\|EF | 3 | 3 | 0 | 8,70E+05 | 24,99 | 2,46E+05 | 95,04 |
| SMARCC2 | sp\|Q8TAQ2\|S | 3 | 3 | 0 | 8,59E+05 | 51,88 | 2,34E+05 | 99,18 |
| RBM27 | sp\|Q9P2N5\|R | 3 | 3 | 0 | 8,38E+05 | 40,46 | 6,23E+05 | 81,61 |
| CSNK2A1 | sp\|P68400\|CS | 3 | 3 | 0 | 8,33E+05 | 50,19 | 3,03E+05 | 70,83 |
| DIDO1 | sp\|Q9BTC0\|D | 3 | 3 | 0 | 8,10E+05 | 62,78 | 4,82E+05 | 109,34 |
| SP1 | sp\|P08047\|SF | 3 | 3 | 0 | 7,17E+05 | 41,97 | 3,67E+05 | 71,53 |
| SMARCA4 | sp\|P51532\|SM | 3 | 3 | 0 | 7,12E+05 | 49,84 | 2,97E+05 | 19,88 |
| ACTL6A | sp\|O96019\|AC | 3 | 3 | 0 | 6,48E+05 | 19,71 | 2,76E+05 | 29,11 |
| CARM1 | sp\|Q86X55\|CA | 3 | 1 | 0 | 6,35E+05 | 122,97 | 5,17E+04 | NA |
| WDR5 | sp\|P61964\|WI | 3 | 3 | 0 | 5,58E+05 | 40,12 | 4,08E+05 | 13,21 |
| PICALM | sp\|Q13492\|PI | 3 | 3 | 0 | 5,45E+05 | 45,03 | 2,28E+05 | 73,76 |
| ANKHD1 | sp\|Q8IWZ3\|AI | 3 | 2 | 0 | 4,76E+05 | 31,00 | 1,86E+05 | NA |
| SAP130 | tr\|A0A2R8YDI | 3 | 2 | 0 | 4,72E+05 | 36,66 | 3,03E+05 | NA |
| FNBP4 | sp\|Q8N3X1\|FI | 3 | 1 | 0 | 4,72E+05 | 14,06 | 1,35E+04 | NA |
| SEC23B | sp\|Q15437\|SC | 3 | 2 | 0 | 4,51E+05 | 56,68 | 6,38E+04 | NA |
| SMARCE1 | sp\|Q969G3\|SI | 3 | 3 | 0 | 4,38E+05 | 42,06 | 1,24E+05 | 101,28 |
| HIVEP1 | sp\|P15822\|ZE | 3 | 3 | 0 | 4,32E+05 | 81,80 | 3,94E+05 | 124,16 |
| SF1 | sp\|Q15637\|SF | 3 | 3 | 0 | 4,18E+05 | 91,49 | 1,06E+05 | 81,67 |
| TCERG1 | sp\|O14776\|TC | 3 | 3 | 0 | 4,17E+05 | 88,58 | 6,51E+05 | 91,24 |
| SEC13 | sp\|P55735\|SE | 3 | 3 | 0 | 4,06E+05 | 65,16 | 1,88E+05 | 44,56 |
| ATF7IP | sp\|Q6VMQ6\|N | 3 | 3 | 0 | 3,68E+05 | 32,98 | 2,39E+05 | 95,57 |
| POU2F1 | sp\|P14859\|PC | 3 | 2 | 0 | 3,45E+05 | 69,32 | 1,44E+05 | NA |
| NCOR1 | sp\|O75376\|NC | 3 | 0 | 0 | 3,29E+05 | 92,78 | 0,00E+00 | NA |
| KANSL3 | sp\|Q9P2N6\|KA | 3 | 3 | 0 | 3,27E+05 | 73,02 | 1,77E+05 | 137,79 |

| | | | | | | | | |
|---|---|---|---|---|---|---|---|---|
| SEC16A | sp\|O15027\|S( | 3 | 1 | 0 | 3,23E+05 | 83,14 | 1,12E+05 | NA |
| ATXN1L | sp\|P0C7T5\|A | 3 | 3 | 0 | 3,06E+05 | 24,45 | 2,69E+05 | 15,06 |
| KPNB1 | sp\|Q14974\|IM | 3 | 3 | 0 | 3,06E+05 | 35,45 | 2,40E+05 | 72,13 |
| NFRKB | sp\|Q6P4R8\|N | 3 | 3 | 0 | 2,91E+05 | 73,62 | 5,60E+04 | 120,51 |
| CSNK2B | sp\|P67870\|CS | 3 | 2 | 0 | 2,91E+05 | 96,46 | 1,05E+05 | NA |
| SMARCC1 | sp\|Q92922\|SN | 3 | 1 | 0 | 2,76E+05 | 33,39 | 9,04E+03 | NA |
| CTTN | sp\|Q14247\|SF | 3 | 3 | 0 | 2,59E+05 | 105,52 | 1,37E+05 | 124,61 |
| SMARCA2 | sp\|P51531\|SN | 3 | 3 | 0 | 2,25E+05 | 24,54 | 1,37E+05 | 77,48 |
| OGT | sp\|O15294\|O( | 3 | 3 | 0 | 2,13E+05 | 58,71 | 1,88E+05 | 58,20 |
| NUP93 | sp\|Q8N1F7\|N | 3 | 3 | 0 | 2,12E+05 | 70,24 | 2,60E+05 | 109,37 |
| TRIM28 | sp\|Q13263\|TI | 3 | 3 | 0 | 2,10E+05 | 60,40 | 2,01E+05 | 87,02 |
| NUP214 | tr\|A0A8Q3SH | 3 | 3 | 0 | 2,01E+05 | 49,31 | 1,65E+05 | 59,99 |
| CLINT1 | sp\|Q14677\|EF | 3 | 0 | 0 | 1,83E+05 | 16,31 | 0,00E+00 | NA |
| RBBP5 | sp\|Q15291\|RE | 3 | 2 | 0 | 1,65E+05 | 67,63 | 2,13E+04 | NA |
| PDLIM5 | sp\|Q96HC4\|P | 3 | 3 | 0 | 1,62E+05 | 31,51 | 1,83E+05 | 63,38 |
| BAP1 | sp\|Q92560\|B/ | 3 | 3 | 0 | 1,61E+05 | 7,60 | 1,86E+05 | 101,89 |
| DMAP1 | sp\|Q9NPF5\|D | 3 | 3 | 0 | 1,61E+05 | 76,74 | 6,92E+04 | 101,65 |
| SP3 | sp\|Q02447\|SF | 3 | 2 | 0 | 1,55E+05 | 137,37 | 1,65E+05 | NA |
| ANXA7 | sp\|P20073\|AN | 3 | 0 | 0 | 1,46E+05 | 43,99 | 0,00E+00 | NA |
| ASH2L | sp\|Q9UBL3\|A | 3 | 2 | 0 | 1,42E+05 | 70,53 | 6,21E+04 | NA |
| SEC24D | sp\|O94855\|S( | 3 | 0 | 0 | 1,39E+05 | 102,35 | 0,00E+00 | NA |
| KIF5B | sp\|P33176\|KI | 3 | 1 | 0 | 1,34E+05 | 67,42 | 3,08E+03 | NA |
| EIF4E | sp\|P06730\|IF4 | 3 | 1 | 0 | 1,29E+05 | 96,65 | 1,59E+04 | NA |
| BCL7C | sp\|Q8WUZ0\|E | 3 | 1 | 0 | 1,26E+05 | 87,37 | 3,01E+04 | NA |
| ENY2 | sp\|Q9NPA8\|E | 3 | 3 | 0 | 1,23E+05 | 21,72 | 4,78E+04 | 80,68 |
| PRPF40A | sp\|O75400\|PF | 3 | 2 | 0 | 1,23E+05 | 45,43 | 1,61E+05 | NA |
| SRF | sp\|P11831\|SF | 3 | 1 | 0 | 1,12E+05 | 73,36 | 3,42E+04 | NA |
| POM121 | sp\|Q96HA1\|P | 3 | 3 | 0 | 1,11E+05 | 55,92 | 8,85E+04 | 119,99 |
| PROSER1 | sp\|Q86XN7\|P | 3 | 2 | 0 | 1,09E+05 | 57,90 | 4,01E+04 | NA |
| SMARCD1 | sp\|Q96GM5\|S | 3 | 1 | 0 | 1,09E+05 | 70,07 | 1,26E+04 | NA |
| ATXN2 | sp\|Q99700\|AT | 3 | 2 | 0 | 1,04E+05 | 117,78 | 3,12E+04 | NA |
| CXXC1 | sp\|Q9P0U4\|C | 3 | 3 | 0 | 1,03E+05 | 76,90 | 2,28E+04 | 41,78 |
| DPY30 | sp\|Q9C005\|DI | 3 | 2 | 0 | 1,01E+05 | 21,74 | 5,50E+04 | NA |
| RC3H2 | sp\|Q9HBD1\|R | 3 | 0 | 0 | 1,01E+05 | 54,45 | 0,00E+00 | NA |
| NRF1 | sp\|Q16656\|NF | 3 | 2 | 0 | 9,97E+04 | 17,21 | 2,98E+04 | NA |
| TBL1XR1 | sp\|Q9BZK7\|TI | 3 | 0 | 0 | 9,62E+04 | 83,33 | 0,00E+00 | NA |
| NOTCH1 | sp\|P46531\|NC | 3 | 0 | 0 | 9,55E+04 | 57,63 | 0,00E+00 | NA |
| DPF2 | sp\|Q92785\|RE | 3 | 1 | 0 | 9,53E+04 | 76,17 | 2,68E+04 | NA |
| VPS72 | sp\|Q15906\|VF | 3 | 2 | 0 | 9,46E+04 | 90,89 | 4,53E+04 | NA |
| AGFG1 | sp\|P52594\|AG | 3 | 0 | 0 | 9,19E+04 | 71,89 | 0,00E+00 | NA |
| ATN1 | sp\|P54259\|AT | 3 | 2 | 0 | 9,02E+04 | 62,15 | 3,75E+04 | NA |
| RRBP1 | sp\|Q9P2E9\|R | 3 | 2 | 0 | 8,96E+04 | 22,74 | 4,20E+04 | NA |
| BICRA | sp\|Q9NZM4\|B | 3 | 0 | 0 | 8,55E+04 | 57,81 | 0,00E+00 | NA |
| SEC24A | sp\|O95486\|S( | 3 | 0 | 0 | 8,17E+04 | 52,33 | 0,00E+00 | NA |
| ILF2 | sp\|Q12905\|ILI | 3 | 1 | 0 | 8,02E+04 | 89,42 | 4,09E+04 | NA |
| POGZ | sp\|Q7Z3K3\|P( | 3 | 2 | 0 | 7,91E+04 | 95,49 | 1,09E+04 | NA |
| SEPTIN2 | sp\|Q15019\|SE | 3 | 2 | 0 | 7,88E+04 | 79,74 | 2,42E+04 | NA |
| RANGAP1 | sp\|P46060\|R/ | 3 | 1 | 0 | 7,44E+04 | 64,47 | 3,15E+03 | NA |
| DDX5 | sp\|P17844\|DE | 3 | 3 | 0 | 7,17E+04 | 86,59 | 8,45E+04 | 69,26 |
| FXR1 | tr\|A0A8V8TM | 3 | 3 | 0 | 7,17E+04 | 69,23 | 7,05E+04 | 33,95 |
| ZC3H10 | sp\|Q96K80\|Z( | 3 | 2 | 0 | 6,79E+04 | 77,79 | 2,56E+04 | NA |
| LSM12 | sp\|Q3MHD2\|L | 3 | 3 | 0 | 6,60E+04 | 76,63 | 3,62E+04 | 48,08 |
| NFYA | sp\|P23511\|NF | 3 | 1 | 0 | 6,05E+04 | 59,48 | 4,22E+04 | NA |
| PPP1R12A | sp\|O14974\|M | 3 | 3 | 0 | 5,76E+04 | 102,67 | 9,07E+04 | 50,57 |
| COPG1 | sp\|Q9Y678\|C( | 3 | 2 | 0 | 5,75E+04 | 122,75 | 3,19E+04 | NA |
| ACTR6 | sp\|Q9GZN1\|A | 3 | 1 | 0 | 5,10E+04 | 49,55 | 4,01E+03 | NA |
| RPN1 | sp\|P04843\|RF | 2 | 3 | 0 | 4,76E+04 | NA | 5,96E+04 | 47,22 |
| TNRC6C | sp\|Q9HCJ0\|TI | 3 | 1 | 0 | 4,59E+04 | 131,32 | 4,55E+04 | NA |

| | | | | | | | | |
|---|---|---|---|---|---|---|---|---|
| ATXN1 | sp\|P54253\|AT | 3 | 2 | 0 | 3,47E+04 | 50,58 | 1,67E+04 | NA |
| CLIC1 | sp\|O00299\|CL | 3 | 2 | 0 | 3,44E+04 | 69,56 | 2,06E+04 | NA |
| PPP2CB | sp\|P62714\|PF | 3 | 1 | 0 | 3,37E+04 | 42,52 | 1,38E+04 | NA |
| FLI1 | sp\|Q01543\|FL | 3 | 0 | 0 | 3,01E+04 | 30,31 | 0,00E+00 | NA |
| CDK1 | sp\|P06493\|CD | 3 | 1 | 0 | 2,69E+04 | 34,16 | 2,52E+04 | NA |
| RBM25 | sp\|P49756\|RE | 3 | 1 | 0 | 2,63E+04 | 66,11 | 6,12E+03 | NA |
| SQOR | sp\|Q9Y6N5\|S | 2 | 3 | 0 | 1,76E+04 | NA | 2,51E+04 | 70,66 |
